# Mapping Spatial Frequency Preferences Across Human Primary Visual Cortex

**DOI:** 10.1101/2021.09.27.462032

**Authors:** William F. Broderick, Eero P. Simoncelli, Jonathan Winawer

## Abstract

Neurons in primate visual cortex (area V1) are tuned for spatial frequency, in a manner that depends on their position in the visual field. Several studies have examined this dependency using fMRI, reporting preferred spatial frequencies (tuning curve peaks) of V1 voxels as a function of eccentricity, but their results differ by as much as two octaves, presumably due to differences in stimuli, measurements, and analysis methodology. Here, we characterize spatial frequency tuning at a millimeter resolution within human primary visual cortex, across stimulus orientation and visual field locations. We measured fMRI responses to a novel set of stimuli, constructed as sinusoidal gratings in log-polar coordinates, which include circular, radial, and spiral geometries. For each individual stimulus, the local spatial frequency varies inversely with eccentricity, and for any given location in the visual field, the full set of stimuli span a broad range of spatial frequencies and orientations. Over the measured range of eccentricities, the preferred spatial frequency is well-fit by a function that varies as the inverse of the eccentricity plus a small constant. We also find small but systematic effects of local stimulus orientation, defined in both absolute coordinates and relative to visual field location. Specifically, peak spatial frequency is higher for pinwheel than annular stimuli and for horizontal than vertical stimuli.

## 1 Introduction

A fundamental goal of visual neuroscience is to quantify the relationship between stimulus properties and neural responses, across the visual field and across visual areas. Studies of primary visual cortex (V1) have been especially fruitful in this regard, with electrophysiological measurements providing good characterizations of the responses of individual neurons to a variety of stimulus properties (Cavanaugh et al., 2002; De Valois et al., 1982; Hubel and Wiesel, 1962; Ringach, 2002). Nearly every neuron in V1 is selective for the local orientation and spatial frequency of visual input, and this has been captured with simple computational models built from oriented bandpass filters (Daugman, 1989; Heeger, 1992; Jones and Palmer, 1987; Pollen and Ronner, 1983; Rust et al., 2005; Vintch et al., 2015).

The characterization of individual neural responses provides only a partial picture of the representation of visual information in V1. In particular, we know that the representation is not homogeneous – receptive field sizes grow and spatial frequency preference decreases with distance from the fovea (eccentricity, De Valois et al., 1982) – but we do not have a general quantitative description of the relationship between these response properties and location in the visual field. There are hundreds of millions of neurons in V1 (Wandell, 1995), and thus, single-unit electrophysiology is unappealing as a methodology for addressing this question.^1^ Functional magnetic resonance imaging (fMRI) offers complementary strengths and weaknesses, allowing simultaneous measurement of responses across all of visual cortex, but at a resolution in which each measurement represents the combined responses of thousands of neurons, limiting the characterization to properties that change smoothly across the cortical surface. Fortuitously, core properties of V1 such as position and spatial frequency tuning *do* vary smoothly across the cortical map (Hubel and Wiesel, 1962; Issa et al., 2000), and so are well suited for summary measures with fMRI. This has led to successful characterization of “population receptive fields” (pRFs), which specify the location and size in visual space of voxel responses (Wandell and Winawer, 2015). A recent study (Aghajari et al., 2020) characterized voxel-wise spatial frequency tuning in early visual cortex, but did not provide an overall description of the dependence of this tuning on retinotopic location or stimulus orientation.

Here, we provide a compact parametric characterization of the spatial frequency and orientation preferences of population receptive fields in area V1, across the visual field. How compact a description can one expect? The information processing of a cortical area such as V1 would be simplest to study and describe if each location in the map analyzed the image with the same computations. This assumption of homogeneous processing is central to signal and image processing, and underlies recent developments in computer vision based on Convolutional Neural Networks (LeCun et al., 1989). But this assumption can be immediately rejected for primate visual systems, since we know that resolution declines precipitously with eccentricity. At the other extreme, if each part of the map analyzed the image in an entirely unique way, the prospect of understanding its function would be hopeless. Fortunately, many properties, such as receptive field size, vary smoothly and systematically with receptive field position, and similar types of models are able to successfully describe neural data across species, individuals, and map locations (e.g., Carandini, 2005).

An attractive intermediate possibility is that cortical processing is conserved across the visual field, up to a dilational scale factor. One hypothesis is that eccentricity-dependent RF scaling emerges first in the RGCs, and then all subsequent stages simply perform a homogeneous (convolutional) transform on their afferents, thus inheriting the eccentricity-scaling of RF sizes. This would result in all neuronal tuning across the cortex being scaled versions of each other. For example, if V1 neurons were tuned such that their preferred spatial frequency was always *p* periods per receptive field, and their receptive fields grew linearly as they moved away from the fovea, such that s = ar (where s is the diameter of the receptive field and *r* is the eccentricity), then neuronal peak spatial frequency would equal *f* = *p/s* = *p/ar*. If this approximates the true relationship between spatial frequency tuning and eccentricity, then sinusoidal gratings, which have a constant frequency everywhere in the image, are an inefficient choice of stimulus to measure this, as high frequencies will be shown at the periphery and low frequencies at the fovea, neither of which will drive responses effectively.

To enable efficient characterization of local spatial frequency preferences, we develop a novel set of global stimuli in which local frequency scales inversely with eccentricity, and which span a variety of orientations. We use these stimuli to probe the dependency of spatial frequency preferences on orientation and retinal location, and summarize this using a compact functional description that is jointly fit to data over the whole visual field. The model parameterization allows spatial frequency tuning to vary with eccentricity, and allows both spatial frequency tuning and BOLD amplitude to vary with retinotopic angle and stimulus orientation. This modeling approach allows flexibility for our parameters of interest, but is not arbitrarily flexible. This is necessary in order to be able to concisely describe how spatial frequency is encoded across the whole visual field and to enable extrapolation to stimuli or visual field positions not included in the study.

## 2 Methods

All experimental materials, data, and code for this project can be found online under the MIT license. Minimally pre-processed data are found on OpenNeuro (Markiewicz et al., 2021), code on GitHub, and other materials on OSF (view README on the GitHub for instructions on how to download and use the data).

### 2.1 Stimulus design

To efficiently estimate preferred spatial frequency across the visual field, we use a novel set of grating stimuli with spatially-varying frequency and orientation. Figure 1 illustrates the logic of the stimulus construction, which is designed for efficient characterization of a system whose preferred spatial frequency falls with eccentricity. Conventional large-field two-dimensional sine gratings will be inefficient for such a system, since the stimulus set will include low-frequency stimuli which are ineffective for the fovea, and high-frequency stimuli which are ineffective for the periphery. Instead, we construct “scaled” log-polar stimuli, such that local spatial frequency decreases in inverse proportion to eccentricity (figure 2B). Specifically, all stimuli are of the form

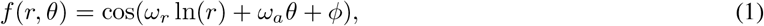

where coordinates (*r, θ*) specify the eccentricity and polar angle of a retinal position, relative to the fovea. The angular frequency *ω_a_* is an integer specifying the number of grating cycles per revolution around the image, while the radial frequency *ω_r_* specifies the number of radians per unit increase in ln(*r*). The parameter *φ* specifies the phase, in radians. The local spatial frequency is equal to the magnitude of the gradient of the argument of cos(·) with respect to retinal position (see Appendix 6.1):

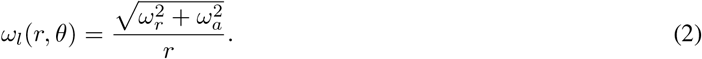

**Figure 1:**
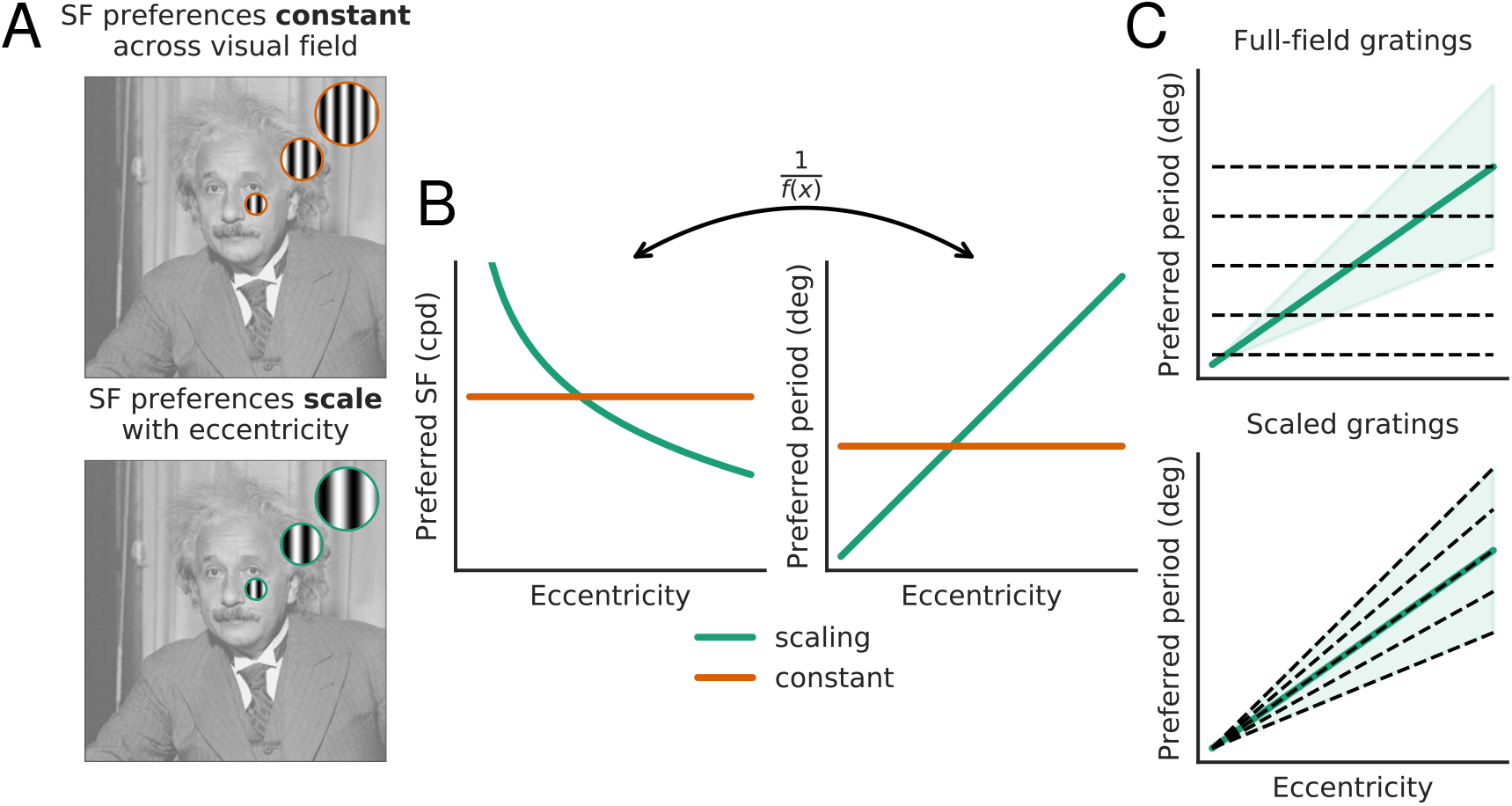
(A) Illustration of two extremal models for spatial frequency preferences across the visual field. Top: preferences are conserved across the visual field (despite changes in receptive field size). Bottom: preferred spatial period (inverse of spatial frequency) is proportional to eccentricity (along with receptive field size). (B) Preferred SF (left) and period (right) as a function of eccentricity, for the two models (red and green curves). (C) Efficiency of stimuli (dashed lines) for probing the scaling model. Top: If preferences scale with eccentricity, conventional full-field two-dimensional sine gratings are an inefficient way to measure spatial frequency tuning: gratings with a large period will be ineffective at driving responses in the fovea and those with a low period will be ineffective for the periphery. Bottom: Oscillating stimuli whose period grows linearly with eccentricity provide a more efficient choice.

**Figure 2:**
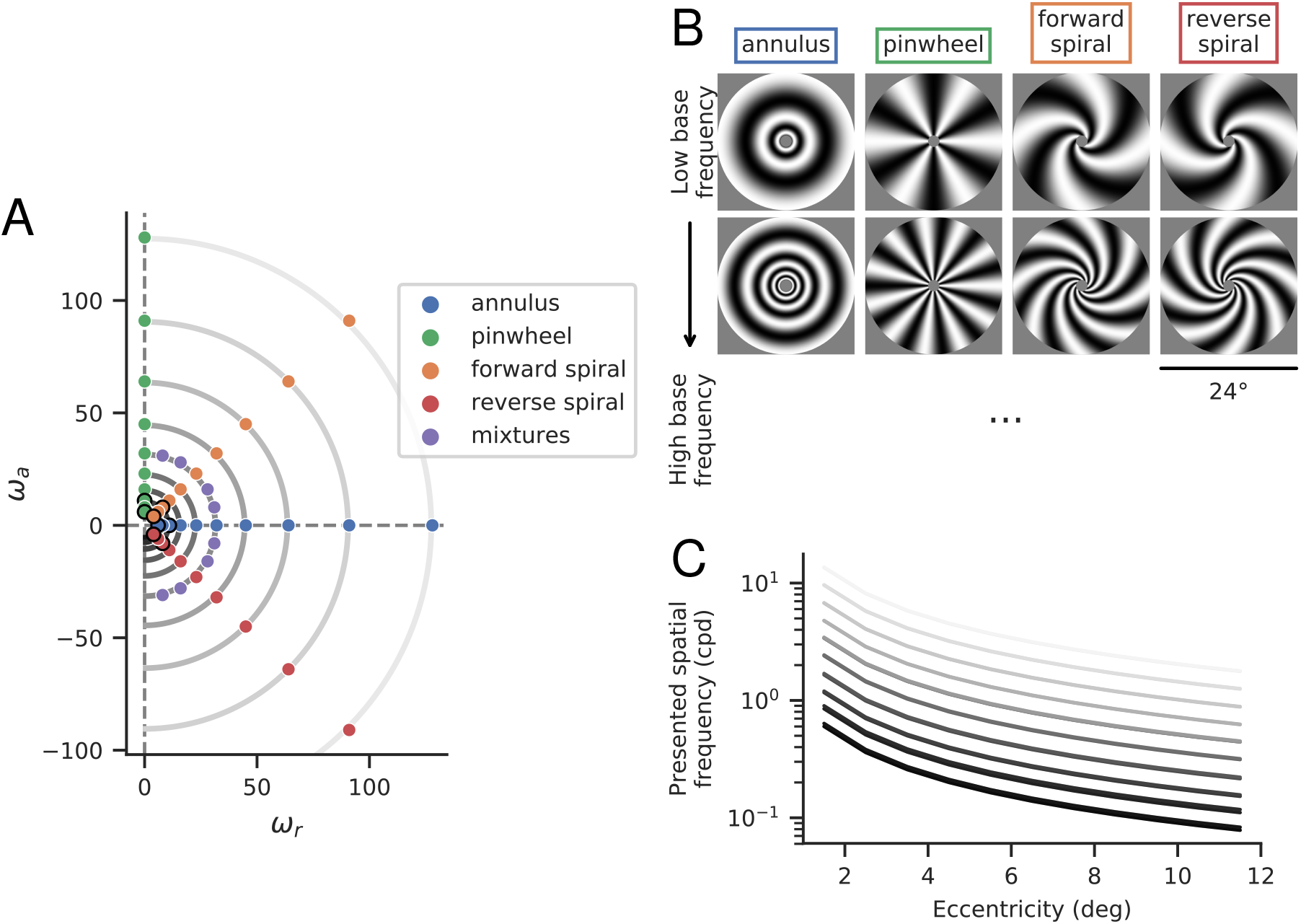
Stimuli. (A) Base frequencies (*ω_r_,ω_a_*) of experimental stimuli. Stimulus category is determined by the relationship between *ω_a_* and *ω_r_*, which determines local orientation information (Eq. 3). (B) Example stimuli from four primary classes, at two different base frequencies. These stimuli correspond to the dots outlined in black in panel A. (C) Local spatial frequencies (in cycles per degree) as a function of eccentricity. Each curve corresponds to stimuli with a specific base frequency, 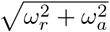, corresponding to the semi-circular contours in panel A. The two rows of stimuli in panel B correspond to the bottom and 3rd-from-bottom curves.

That is, local frequency is equal to Euclidean norm of the frequency vector (*ω_r_, ω_a_*) divided by eccentricity (in units of radians per pixel), which implies that the local spatial period of the stimuli grows linearly with eccentricity. Similarly, the local orientation can be obtained by taking the angle of the gradient of the argument of cos(·) with respect to retinal position (see Appendix 6.1):

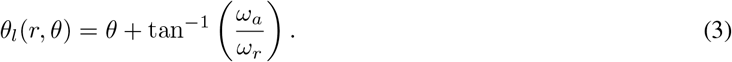

That is, the local grating orientation is the angular position relative to the fovea, plus the angle of the two-dimensional frequency vector (*ω_r_, ω_α_*). Note that *θ_ι_* is in absolute units (e.g., *θ_ι_* =0 indicates local orientation is vertical, regardless of location). For our stimuli, this depends on the polar angle, but a uniform grating has the same *θ_ι_* value everywhere in the image (its orientation thus does not depend on polar angle).

We generated stimuli corresponding to 48 different frequency vectors (see Fig. 2), at 8 different phases *ϕ* ∈ {0,*π*/4,*π*/2,..., 7*π*/4}. The frequency vectors were organized into five different categories:

1. Pinwheels: *ω_r_* = 0, *ω_a_* ∈ {6; 8; 11; 16; 23; 32; 45; 64; 91; 128}
2. Annuli: *ω_a_* = 0, *ω_r_* ∈ {6; 8; 11; 16; 23; 32; 45; 64; 91; 128}
3. Forward spirals: *ω_r_* = *ω_a_* ∈ {4; 6; 8; 11; 16; 23; 32; 45; 64; 91}
4. Reverse spirals: *ω_r_* = −*ω_a_* ∈ {4; 6; 8; 11; 16; 23; 32; 45; 64; 91}
5. Fixed-frequency mixtures: (*ω_r_*; *ω_a_*) ∈ {(8; 31); (16; 28); (28; 16); (31; 8); (31; –8); (28; –16); (16; –28); (8; –31)}

Note that *ω_a_* values must be integers (since they specify cycles per revolution around the image), and we chose matching integer values for *ω_r_*. Because of this constraint, the pinwheel/annulus and the forward/reverse spiral stimuli have slightly different local spatial frequencies. For the same reason, the local spatial frequency of the mixture stimuli is only approximately matched across stimuli 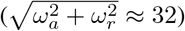. Across all stimuli, the spatial frequencies presented at any given eccentricity span a 20-fold range (figure 2C). For example, at the most foveal portion of the stimuli (from 1 to 2 deg) the frequencies are log-spaced from 0.6 to 13.65 cpd. In the most peripheral region (11 to 12 deg)), the range is 0.078 to 1.78 cpd.

### 2.2 Display Calibration

The projector used to display stimuli in our experiments was calibrated to produce light intensities proportional to luminance. In addition, we wanted to compensate for spatial blur (due to a combination of display electronics or optics) that could systematically alter the frequency content of our stimuli. We estimated the modulation transfer function (MTF) of the projector (i.e., the Michelson contrast as a function of spatial frequency), shown in figure 3. We used a calibrated camera and developed custom software to process and analyze photographs of full-contrast square-wave gratings. We found that the contrast of the projected image decreased by roughly 50% as it approached the Nyquist frequency of 0.5 cycles per display pixel. We compensated for these effects by rescaling the amplitude of low frequency content in our stimuli, by an amount proportional to the inverse MTF (note that the more natural procedure of increasing the high frequency content is not practical, as it could exceed the maximum contrast that can be displayed).

**Figure 3:**
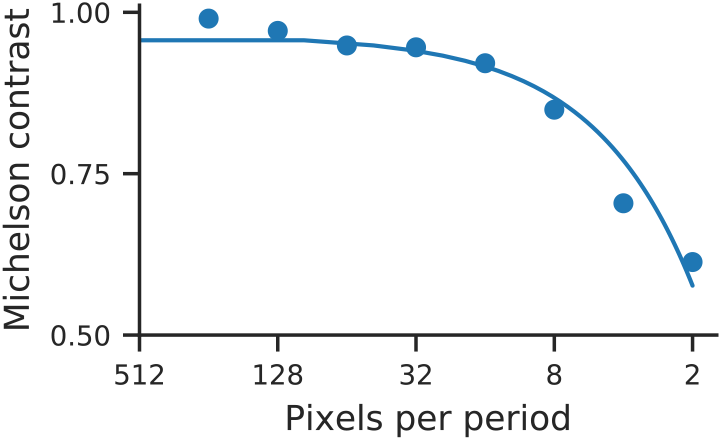
Estimated modulation transfer function (MTF) of the projector used in our experiments. Michelson contrast was measured for periods from 2 to 256 pixels (blue points) and then fit with a univariate spline (blue curve) with smoothing degree 1 (Virtanen et al., 2020). The fitted spline was used for calibration.

### 2.3 Participants

Twelve participants (7 women and 5 men, aged 22 to 35), including an author (W.F.B.), participated in the study and were recruited from New York University. All subjects had normal or corrected-to-normal vision. Each subject completed 12 runs, except for sub-04, who only completed 7 of the 12 runs due to technical issues that came up during the run. The quality of their GLMdenoise fits and their final model fits do not appear to vary much from that of the other subjects. All subjects provided informed consent before participating in the study. The experiment was conducted in accordance with the Declaration of Helsinki and was approved by the New York university ethics committee on activities involving human subjects.

### 2.4 Experimental Design

The experiment was run on an Apple MacIntosh computer, using custom scripts with PsychoPy (Peirce et al., 2019a), presented on a luminance-calibrated MTF-corrected VPixx ProPixx projector. Images were projected onto a screen, which the subject viewed through a mirror. The screen was 36.2 cm high and 83.5 cm from the subject’s eyes (73.5 cm from screen to mirror, and approximately 10 cm from mirror to eyes). Stimuli were constrained to a circular aperture filling the height of the display (12 deg radius), with an anti-aliasing mask at the center (0.96 deg radius). Each stimulus class was presented in a 4-s trial, during which the 8 images with different phases were shown in randomized order. Each of the 8 images was presented once, cycled on and off (300-msec on, 200-msec off) in order to minimize adaptation. A movie of a single run can be viewed on the OSF. Each of the 48 stimulus classes was presented once in each of 12 runs, with the presentation order of the stimulus classes and of the phases randomized across runs. Subjects viewed these stimuli while performing a one-back task on a stream of alternating black and white digits (1-sec on, 1-sec off) at the center of the screen in order to ensure accurate fixation, minimize attentional effects, and maintain a constant cognitive state. Thus, the central one degree of vision always contained either a blank midgrey screen or a black or white digit. This lessens the possibility of differences in fixational eye movements that might arise from differences in stimulus structure near the fovea. Behavioral responses were recorded using a button box (see appendix 6.2 for behavioral analysis).

### 2.5 fMRI Scanning Protocol

All MRI data for the spatial frequency experiment were acquired at the NYU Center for Brain Imaging using a 3T Siemens Prisma scanner with a Siemens 64 channel head/neck coil. For fMRI scans, we used the CMRR MultiBand Accelerated EPI Pulse Sequence (Release R015a) (TR, 1000 ms; TE, 37 ms; voxel size, 2mm^3^; flip angle, 68 degrees; multiband acceleration factor, 6; phase-encoding, posterior-anterior) (Feinberg et al., 2010; Moeller et al., 2010; Xu et al., 2013). High resolution whole-brain anatomical T1-weighted images (1 mm^3^ isotropic voxels) were acquired from each subject for registration and segmentation using a 3D rapid gradient echo sequence (MPRAGE). Two additional scans were collected with reversed phase-encoded blips, resulting in spatial distortions in opposite directions. These scans were used to estimate and correct for spatial distortions in the EPI runs using a method similar to Andersson et al., 2003, as implemented in FSL (Smith et al., 2004).

### 2.6 Preprocessing

fMRI data were minimally preprocessed using a custom script (available on the Winawer lab Github) which builds a Nipype (Gorgolewski et al., 2018; Gorgolewski et al., 2011) pipeline. Brain surfaces were reconstructed using `recon-all` from FreeSurfer v6.0.0 Dale et al., 1999. Functional images were motion corrected using `mcflirt` (FSL v5.0.10 Jenkinson et al., 2002) to the single-band reference image gathered for each scan. Each single-band reference image was then registered to the distortion scan with the same phase-encoding direction using `flirt` (FSL v5.0.10 Greve and Fischl, 2009; Jenkinson et al., 2002; Jenkinson and Smith, 2001). Distortion correction was performed using an implementation of the TOPUP technique Andersson et al., 2003 using TOPUP and `ApplyTOPUP` (FSL v5.0.10 Smith et al., 2004). The unwarped distortion scan was co-registered to the corresponding T1w using boundary-based registration Greve and Fischl, 2009 with 9 degrees of freedom, using `bbregister` (FreeSurfer v6.0.0). The motion correcting transformations and BOLD-to-T1w transformation were concatenated using `ConvertXFM` (FSL v5.0.10) and then were applied to the functional runs in a single step along with the unwarping warpfields using `ApplyWarp` (FSL v5.0.10). Applying the corrections in a single step minimizes blurring from the multiple interpolations.

### 2.7 Retinotopy

A separate retinotopy experiment was used to obtain the population receptive field (pRF) location and size for V1 voxels in each subject (Wandell and Winawer, 2015). This experiment consisted of six standard pRF mapping runs, with sweeping bar contrast apertures filled with a variety of colorful objects, faces and textures. This stimulus has been shown to be effective in evoking BOLD responses across many of the retinotopic maps in visual cortices (Benson and Winawer, 2018; Benson et al., 2018; Himmelberg et al., 2021). The results of this pRF mapping were combined with a retinotopic atlas (Benson et al., 2014) in order to improve the accuracy of the retinotopic map (see Benson and Winawer, 2018 for a description of this method). The stimulus, fMRI acquisition parameters, and fMRI pre-processing for the retinotopy experiments are described in detail in Benson and Winawer, 2018 and Himmelberg et al., 2021.

### 2.8 Stimulus response estimation

Response amplitudes were estimated using the GLMdenoise MATLAB toolbox (Kay et al., 2013a). The algorithm fits an observer-specific hemodynamic response function (HRF), estimating response amplitudes (in units of percent BOLD signal change) for each voxel and for each stimulus, with 100 bootstraps across runs. Thus for each voxel we estimate 48 responses (one for each unique pair (*ω_α_, ω_r_*), averaged over the 8 phases shown within the trials). The algorithm also includes three polynomial regressors (degrees 0 through 2) to capture the signal mean and slow drift, and noise regressors derived from brain voxels that are not well fit by the GLM.

The combined retinotopy and GLMdenoise measurements consist of (for each voxel): the visual area, population receptive field location and size, and 100 bootstrapped response amplitudes to each of the 48 stimuli.

### 2.9 One-Dimensional Tuning Curves

We fit one-dimensional log-normal tuning curves to the responses of groups of voxels at different eccentricities (lying within one-degree eccentricity bins):

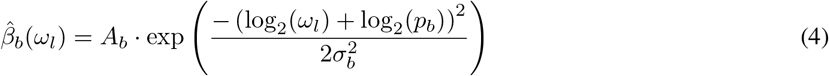

where 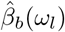 is the average BOLD response in eccentricity bin *b* at spatial frequency *ω_ι_* (in cycles per degree), *A_b_* is the response gain, *p_b_* is the preferred period (the reciprocal of the peak spatial frequency, *ω_b_*, which is the mode of the tuning curve), and *σ_b_* is the bandwidth, in octaves. Fits were obtained separately for the four primary stimulus classes (pinwheel, annulus, forward spiral, and reverse spiral).

We fit these tuning curves 100 times per subject, per stimulus class, and per eccentricity, bootstrapping across the fMRI runs (12 per subject).

### 2.10 Two-Dimensional Tuning Curves

Our one-dimensional tuning curves are averaged over stimulus orientation and retinotopic angle. To capture the effect of these additional stimulus attributes, we developed a two-dimensional model for individual voxel responses as a function of stimulus local spatial frequency (in cycles per degree), *ω_ι_*, stimulus local orientation, *θ_ι_*, voxel eccentricity (in degrees), *r_v_*, and voxel retinotopic angle, *θ_v_* (figure 4A). Responses are again assumed to be log-normal with respect to spatial frequency:

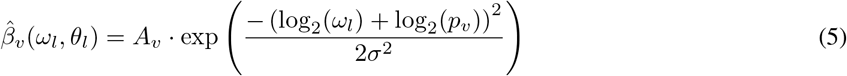

**Figure 4:**
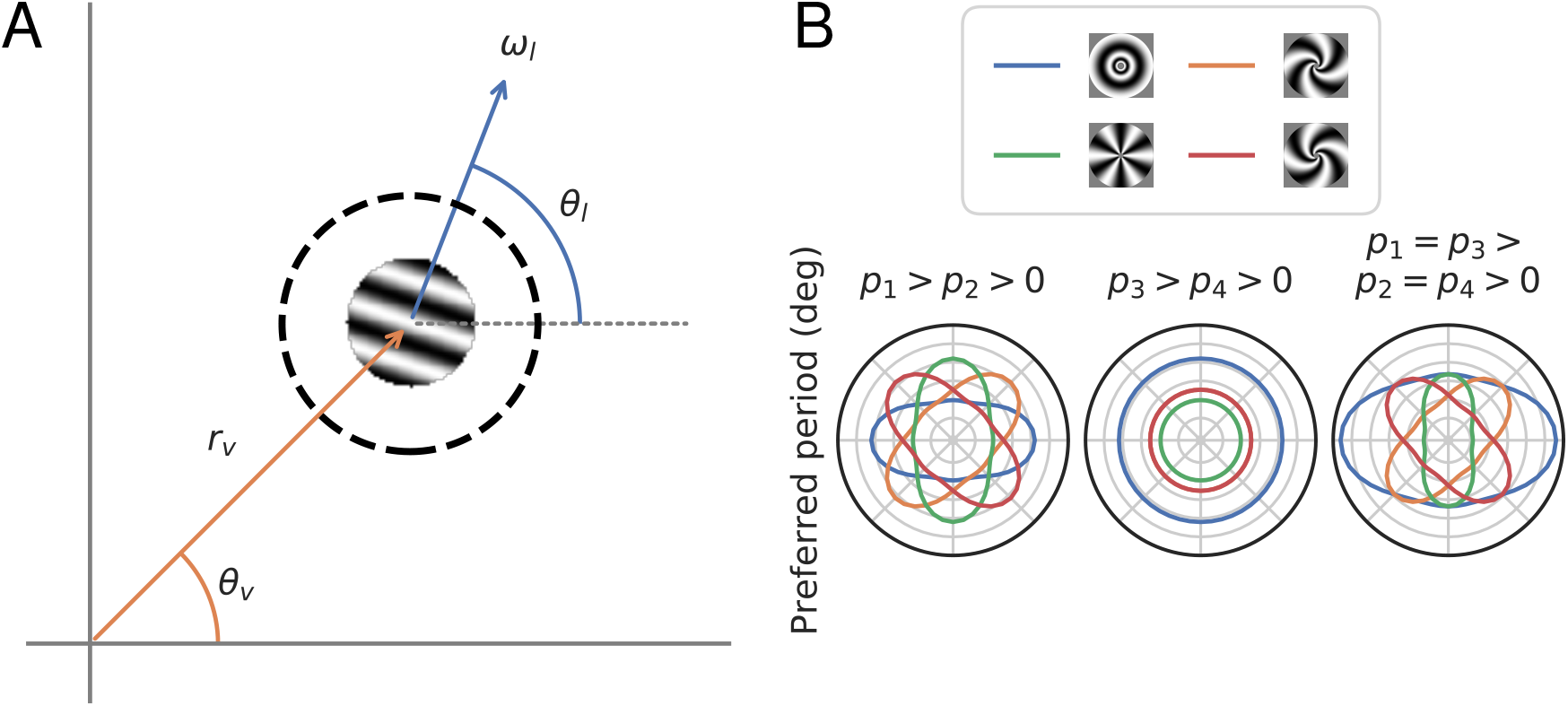
(A) Local stimulus parameterization for the two-dimensional model. The model is a function of four variables, two related to voxel population receptive field location and two related to stimulus properties. *r_v_* and *θ_v_* specify the eccentricity (in degrees) and the retinotopic angle of the location of the center of the voxel’s population receptive field, relative to the fovea. *ω_ι_* and *θ_ι_*, specify the local spatial frequency (in cycles per degree) and the local orientation (in radians, counter-clockwise relative to horizontal) of the stimulus, at the center of that voxel’s population receptive field (dashed line). (B) Schematic showing the effects of *p_i_* parameters on preferred period as a function of retinotopic angle at a single eccentricity for the four main stimulus types used in this experiment. When *p*_1_ > *p*_2_ > 0 (and *p*_3_ = *p*_4_ = 0), the effect of orientation on preferred period is in the absolute reference frame only, i.e., preferred period only depends on absolute orientation (e.g., vertical or horizontal). In this plot, preferred period varies with retinotopic angle because the absolute orientation of our stimuli vary with retinotopic angle (for another example, see fig 10, where the relative amplitude effect is also only in the absolute reference frame; thus the relative amplitude is always higher for vertical than horizontal stimuli). When *p*_3_ > *p*_4_ > 0 (and *p*_1_ = *p*_2_ = 0), the effect of orientation is in the relative reference frame only, and annulus stimuli will always have the highest preferred period. Finally, when all *p_i_* = 0, the effects are mixed.

In our one-dimensional analysis, we fit parameters {*p, A, σ*} separately to each eccentricity band and stimulus class. Based on the results of that analysis (see 3.1), we assume *σ* is constant across eccentricities, retinal position, and local stimulus spatial frequency (while others have found some variation in bandwidth with respect to these variables, this study focuses on peak spatial frequency tuning and we do not include extra flexibilty in model bandwidth, in order to avoid overfitting). We assume functional forms for the dependencies of parameters *p* and *A* on retinal position, local stimulus spatial frequency, and local stimulus orientation. First, we parameterize the effect of eccentricity, fitting the preferred period as an affine function of a voxel’s eccentricity *r_v_: p_v_* = *ar_v_* + *b*. We assume that this baseline dependency is modulated by effects of retinotopic angle and stimulus orientation, both of which are known to affect visual perception (Barbot et al., 2020; Heeley and Timney, 1988; Williams et al., 1981). Specifically, we express preferred period as:

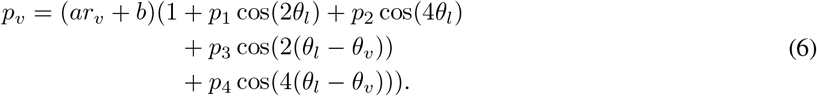

The parameters *p_i_* have the following interpretations:

*p*_1_: absolute cardinal effect, horizontal vs. vertical. A positive pi means that voxels have a higher preferred period for vertical than for horizontal stimuli.
*p*_2_: absolute cardinals vs. obliques effect, horizontal/vertical vs. diagonals. A positive *p*_2_ means that voxels have a higher preferred period for cardinal than for oblique stimuli.
*p*_3_: relative cardinal effect, annuli vs. pinwheels. A positive *p*_3_ means that voxels have a higher preferred period for annular than for pinwheel stimuli.
*p*_4_: relative cardinals vs. obliques effect, annuli/pinwheels vs. spirals. A positive *p*_4_ means that voxels have a higher preferred period for annuli and pinwheels than for spirals.

*p*_1_ and *p*_2_ have effects in the absolute reference frame because they only depend on *θ_ι_*, the orientation in absolute terms, whereas *p*_3_ and *p*_4_ additionally depend on *θ_v_* and thus have effects in the relative reference frame.

To illustrate these effects, we show tuning functions for several stimulus classes given a few possible parameter combinations (figure 4B). We also provide an interactive tool that enables the user to set arbitrary values for all parameters and to probe how the parameter settings influence the pattern of responses to various stimulus types (this website).

We also express the gain of the BOLD responses as a function of voxel retinotopic angle and stimulus orientation (without the eccentricity-dependent base term):

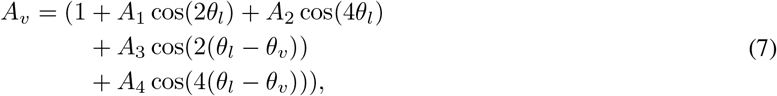

This parameterization allows the amplitude to vary depending on both absolute stimulus orientation (*θ_ι_*), and stimulus orientation relative to retinotopic angle (*θ_ι_ – θ_v_*), but not on absolute retinotopic location. This choice is premised on the fact that voxel-to-voxel variation in the amplitude of the BOLD signal depends in part on factors that are not neural and are not of interest here. For example, BOLD amplitude is influenced by draining veins (Kay et al., 2019; Lee et al., 1995) and the orientation of the gray matter surface relative to the instrument magnetic field (Gagnon et al., 2015), as well as other factors not directly related to neural responses.

In addition, the model cannot capture categorical differences across the visual field, e.g., between upper and lower, or foveal and parafoveal visual field, except insofar as the parametric forms allow (linear function of eccentricity, harmonics of stimulus orientation).

### 2.11 Model fitting

We fit the 2D model to all V1 voxels simultaneously, excluding voxels whose population receptive field (pRF) center lies outside the stimulus, those whose pRF center lies within one standard deviation of the stimulus border, and those with an average negative response to our stimuli. Voxels with negative responses but whose pRFs are centered within the stimulus extent are likely dominated by artifacts such as those arising from draining veins (Lee et al., 1995; Winawer et al., 2010).

The remaining voxels vary widely in their signal to noise ratio. Typically in fMRI analyses, all voxels whose noise level lies above some threshold are excluded from the analysis. Here, we instead weight each voxels’ loss by its precision, so that noisier voxels will contribute less to the parameter estimates. Specifically, we use a normalized mean-squared error loss over voxels:

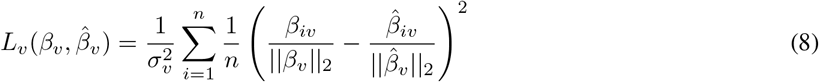

where *i* indexes the *n* different stimulus classes, *β_iv_* is the response of voxel *v* (estimated using GLMdenoise) to stimulus class *i*, 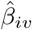 is the response to stimulus class *i* predicted by our model, ||*β_v_*||_2_ is the L2-norm of *β_v_* (across all stimulus classes), and 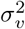 is the variance of voxel *v*’s response (that is, 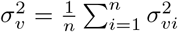, where *σ_vi_* is half of the 68 percentile range of the response of voxel *v* to stimulus class *i*, as estimated by GLMdenoise). This loss function is equivalent to the cosine between response vectors *β_v_* and 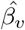 multiplied by 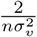. Normalization of the *β_v_* and 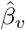 vectors allows the fitting to be agnostic to variations in absolute response amplitude, capturing the response dependency on stimulus and retinal location.

We minimize the average of this loss across all appropriate voxels, using custom code written in PyTorch (Paszke et al., 2019) and using the AMSGrad variant of the Adam optimization algorithm (Kingma and Ba, 2014; Reddi et al., 2018). To assess model accuracy, we use 12-fold cross-validation (see 3.2.1). Specifically, we fit the model to 44 of the 48 stimulus classes, then get predictions for the 4 held-out classes. We do this for each of the 12 subsets, which get us a complete 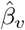 that we can compare against *β_v_*.

### 2.12 Software

Data analysis, modeling, and figure creation were done using a variety of custom scripts written in Python 3.6.3 (Van Rossum and Drake, 2009), all found in the GitHub repository associated with this paper. The following packages were used: snakemake (Mölder et al., 2021), Jupyter Lab (Kluyver et al., 2016), numpy (“Array programming with NumPy”, 2020), matplotlib (Hunter, 2007), scipy (Virtanen et al., 2020), seaborn (Waskom, 2021), pandas (McKinney, 2010; pandas development team, 2020), nipype (Gorgolewski et al., 2018; Gorgolewski et al., 2011), nibabel (Brett et al., 2020), scikit-learn (Pedregosa et al., 2011), neuropythy (Benson and Winawer, 2018), pytorch (Paszke et al., 2019), psychopy (Peirce et al., 2019b), FSL (Smith et al., 2004), freesurfer (Dale et al., 1999), vistasoft, and GLMdenoise (Kay et al., 2013a).

## 3 Results

### 3.1 One-Dimensional Analysis

We start by analyzing the data as a function of spatial frequency alone (i.e., averaging over orientation), which requires fewer assumptions and is easier to visualize. We fit log-normal tuning curves to averaged voxel responses at each eccentricity for each of the four main stimulus classes. The log-normal function provides a reasonably good fit to the data (see figure 5).

**Figure 5:**
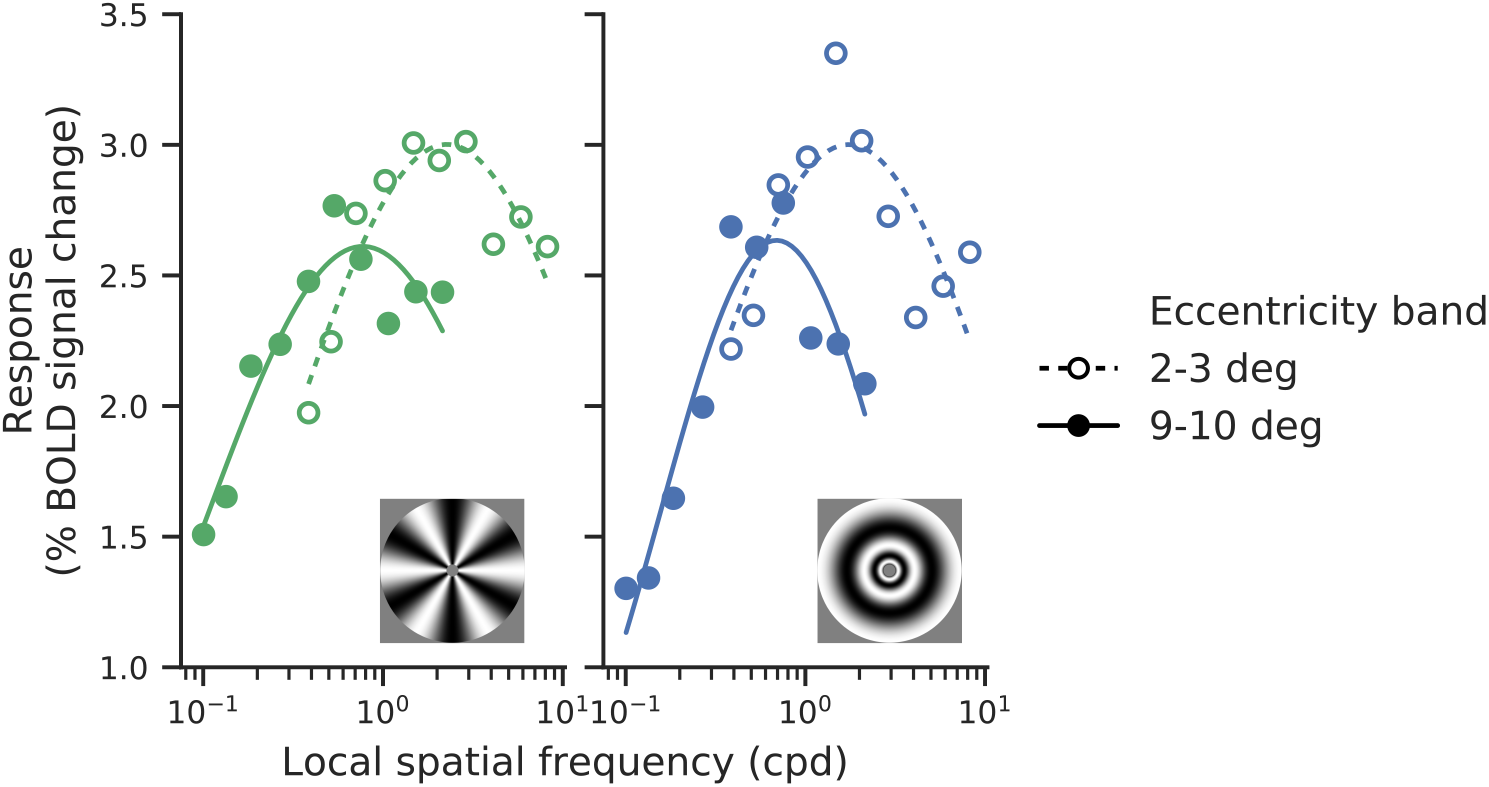
Example data and best-fitting log-normal tuning curves for responses of one subject (sub-01) to pinwheel (left) and annular (right) stimuli. The solid line and filled circles correspond to 9-10deg eccentricity, while dashed line and empty circles correspond to 2-3deg.

We then combined the preferred periods across subjects by bootstrapping a precision-weighted mean: for each eccentricity and stimulus class, we selected 12 subjects at random with replacement, multiplied each subject’s median preferred period by the precision of that estimate, and averaged the resulting values:

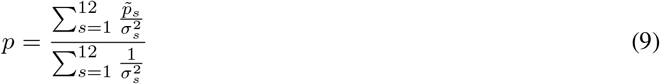

where 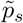 is the median preferred period value for subject *s* and *σ_s_* is the difference between the 16th and 84th percentile for that subject. This bootstrapping is done 100 times to obtain median values and 68% confidence intervals displayed in figure 6A. The precision weighted average has the virtue of giving more weight to better parameter estimates while not fully discarding data.

**Figure 6:**
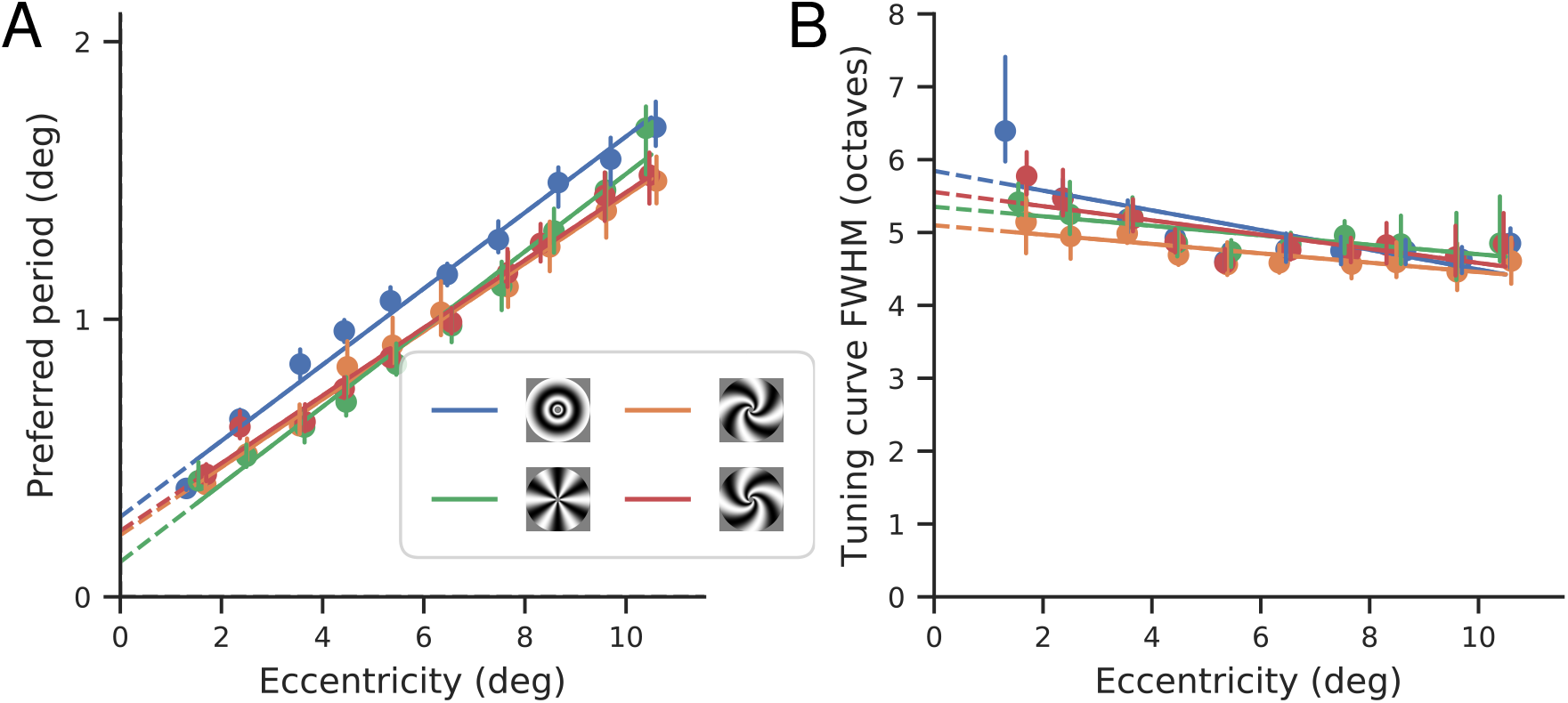
Spatial frequency tuning. (A) Preferred period of tuning curves (parameter *p_b_* in equation (9), *n* = 12), as functions of eccentricity, fit separately for the four different stimulus classes. Points and vertical bars indicate the median and 68% confidence intervals obtained from bootstraps combining subjects using a precision-weighted average (see text). Lines are the best linear fits. (B) Full-width half-maximum (in octaves) of tuning curves, as functions of eccentricity, fit separately for the four different stimulus classes. Points and vertical bars indicate the median and 68% confidence intervals obtained from bootstraps combining subjects using a precision-weighted average (see text). Lines are the best linear fits.

The preferred period for each stimulus class is well-described as an affine function of eccentricity, with a positive offset. Thus, the spatial frequency preferences of V1 do not scale perfectly with eccentricity (e.g., the preferred frequency at 4 degrees is not half that of 2 degrees). There is also a noticeable dependence on stimulus orientation, with the annular stimuli exhibiting a larger preferred period than the other three stimuli at each eccentricity. Differences between the other stimulus types are more subtle, but perhaps indicate a slightly reduced slope for the two spiral stimuli relative to the pinwheel.

We do the same precision-weighted bootstrapping process for the full-width half-maximum (in octaves) of the tuning curves shown in 6B. We can see that the FWHM is mostly constant across eccentricities, except for some larger, noisier values for the most foveal voxels. We believe this apparent dip is due to how the fits are constrained, rather than a real decline in tuning curve width: as can be seen in figure 8A, the presented frequencies shift from the right of the tuning curve to the left for more peripheral voxels. In the periphery and the fovea, where most of the presented frequencies fall on one side of the curve, the width is unlikely to be well-constrained, resulting in the higher error bars seen in figure 6B. FWHM additionally appears to be consistent across stimulus types.

### 3.2 Two-Dimensional Model

The one-dimensional model provides a useful but limited overview of spatial frequency selectivity. In particular, we’ve treated the four stimulus classes as discrete categories, rather than members of a continuum over relative orientation. Moreover, this analysis conflates the effects of absolute orientation (relative to a global vertical/horizontal coordinate system) and orientation relative to a voxel’s retinotopic angle. These might be systematically different, and because there are more voxels at some retinotopic angles than others (e.g., Benson et al., 2020; Silva et al., 2018), the averaging might cause systematic biases in the summary measures. Finally, the analysis examines peak spatial frequency tuning but does not examine possible differences in BOLD amplitude for different stimulus orientations.

The two-dimensional model described in section 2.10 allows us to more directly and comprehensively assess how spatial frequency tuning varies across the visual field. Instead of binning voxels by eccentricity, we fit all voxels simultaneously, with each voxel’s contribution to the loss function weighted by the precision of its responses. By fitting each voxel, we can tease apart the effects of absolute and relative orientation (figure 4B). We are able to parameterize these effects on both preferred period and gain. Finally, the fitted model will generate predictions for the response of any voxel in the visual field to any spatial frequency and orientation (though its predictions will likely decrease in accuracy the farther the voxel’s retinotopic location and stimulus properties move from the those included in this study).

#### 3.2.1 Model selection

The full 2D model has 11 parameters, and we used cross-validation in order to determine which are necessary to explain the data in V1. Omitting or including all combinations of parameters would yield 2^11^ possible models. To reduce this, we grouped the parameters into several small sets, based on whether they affect the preferred period or gain and whether their effect is determined by eccentricity, relative orientation, or absolute orientation. For example, *p*_1_ and *p*_2_ both affect preferred period as a function of absolute orientation and so are always both present or both absent. Moreover, we only tested parameter combinations that we considered plausible; for example, we do not test relative preferred period and absolute gain. Figure 7A shows the 14 candidate submodels considered. When fitting model 8, for example, the parameters *σ, a, b,p*_1_,*p*_2_, *A*_1_, *A*_2_ are all fit, while *p*_3_,*p*_4_, *A*_3_, *A*_4_ are set to 0; this corresponds to modeling the preferred period as a linear function of eccentricity, modulated by absolute orientation, and modeling the gain as also modulated by absolute orientation.

**Figure 7:**
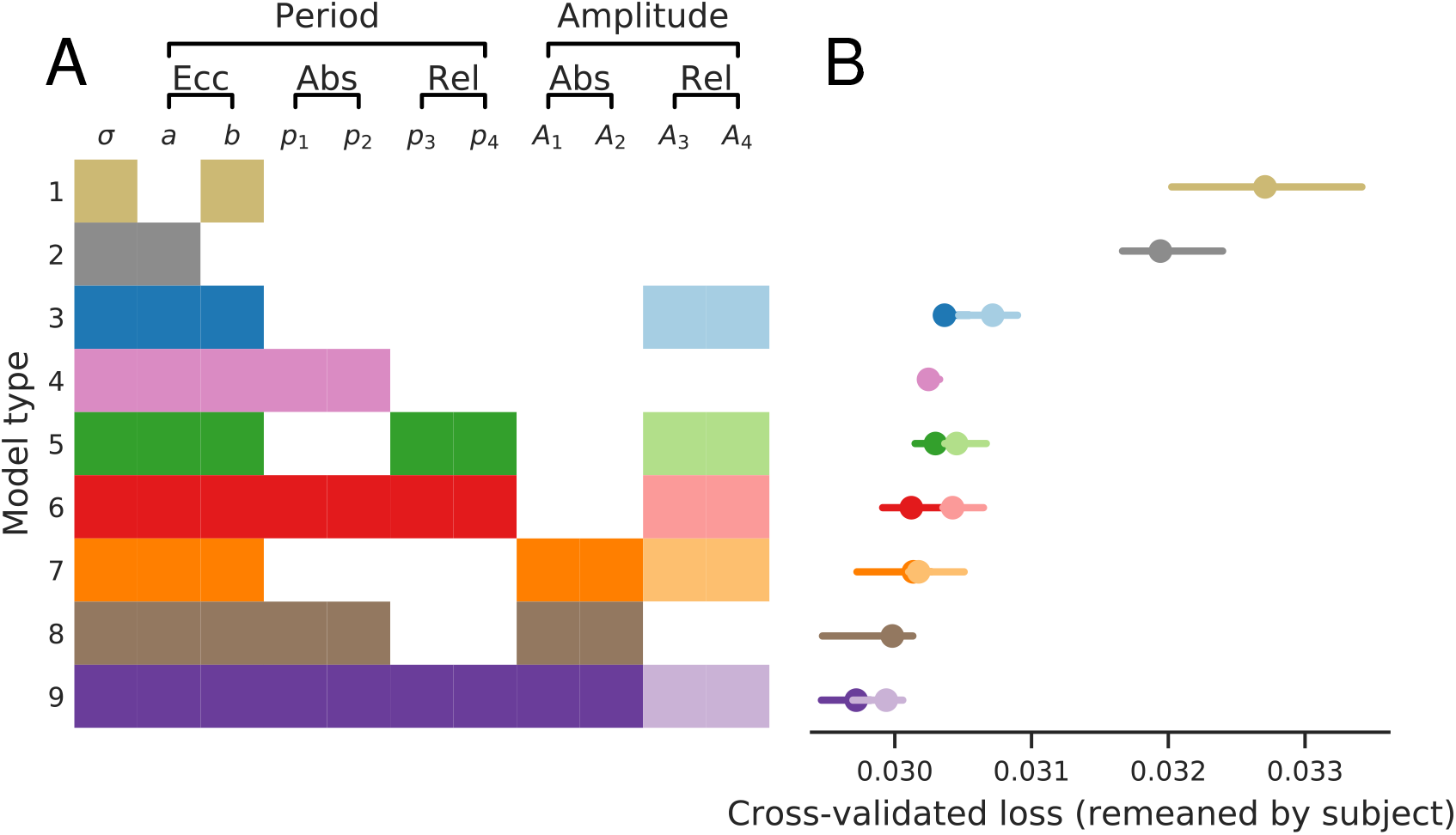
Nested model comparison via cross-validation. (A) 14 different submodels are compared to determine which of the 11 parameters, as defined in Equations (4), (6), and (7), are necessary. Model parameters are grouped by whether they affect the period or the gain, and whether their effect relates to eccentricity, absolute orientation, or relative orientation. Filled color boxes indicate parameter subset used for each submodel. (B) Cross-validated loss for each submodel. Models are fit to each subject separately, using 12-fold cross-validation (each fold leaves out 4 random stimuli). Quality of fit varies across subjects, so to combine subjects and view the effect of model, we subtract each subject’s mean loss across models, then add back the average loss across subjects and models. Bars show the 68% confidence intervals from bootstrapped mean across subjects.

Submodels are fit per subject, with 12-fold cross-validation, withholding four random stimuli from fitting on each fold, using the same partitions across models and subjects. After training, predictions are generated for these 4 stimuli, and the subject’s cross-validation loss for the model is computed across all of the held-out data (12 folds). Cross-validation loss varies greatly across subjects, dependent on the subject’s signal to noise ratio. To combine across subjects, we normalize the data by subtracting each subject’s mean cross-validation loss across models. For visualization, we then add back the average loss across subjects. Figure 7B shows the median cross-validation loss and 68% confidence intervals of these losses. For some rows, two models are shown: the model with and without fitting parameters *A*_3_, *A*_4_. (The variant that fits those parameters is shown in the desaturated color.) The results indicate that 9 of the 11 parameters contribute to accurately predicting responses. By fitting each of the 14 candidate models to each subject individually, we find that all parameter groupings improve performance except for *A*_3_ and *A*_4_: the loss is greater whenever those two are included.

Comparing the losses of models 1, 2, and 3 reveals the importance of the two parameters relating eccentricity to preferred period: while a line through the origin (model 2) captures the data better than a constant value (model 1), the performance increases substantially with an affine model using both terms (model 3). In sum, both parameters *a* and b are required to accurately explain the data, and preferred period increases linearly with eccentricity with a non-zero intercept.

Beyond eccentricity, the effect of orientation on preferred period does not change performance much unless one also adds the effect on gain (models 4 through 6 all have similar performance). The effect of relative orientation on gain by itself has a negative effect on performance, as can be seen by comparing the saturated and desaturated points for models 3, 5, 6, 7, and 9. Absolute orientation, on the other hand, improves performance, as can be seen by comparing 6 and 9, 4 and 8, or 3 and 7. Therefore, for the remainder of this paper, we use the saturated point of model 9, which has the lowest cross-validation loss and fits all preferred period parameters, *p_k_*, as well as those that capture the effect of absolute orientation on gain.

#### 3.2.2 Spatial frequency tuning across stimulus orientation and visual field positions

Having selected model 9, we then re-fit it to each subject without cross-validation. Specifically, we fit model 9 to each of 100 bootstraps from each subject separately, giving us 100 estimates of each model parameter per subject. Figure 8A shows three example voxels’ median responses and model 9’s median predictions, as a function of local spatial frequency, from one subject. As expected, the peak of the spatial frequency tuning function decreases with increasing eccentricity. The bandwidth (in octaves) is comparable across eccentricities, and the plots indicate that the stimuli sampled the local spatial frequencies appropriately at each eccentricity.

**Figure 8:**
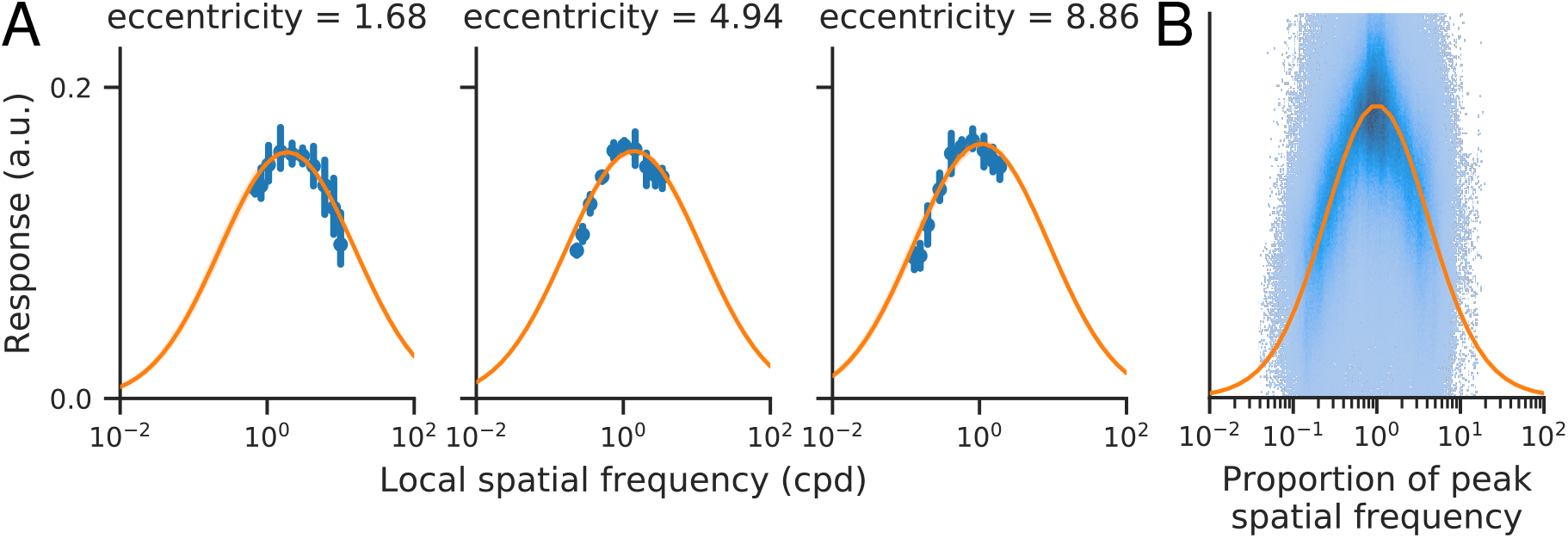
(A) Three example voxels from a single subject (sub-01). Blue points indicate median voxel responses across bootstraps. Error bars indicate variation as a function of orientation. Orange line shows model 9’s predictions, in both cases as a function of the local spatial frequency at the center of each voxel’s pRF. (B) Responses of all voxels across all subjects as two-dimensional histogram. For each voxel and stimulus orientation, responses are plotted as a function of spatial frequency, relative to peak spatial frequency. Orange line shows model 9’s predictions.

Overall, the log-Gaussian tuning function provided a good fit to the complete dataset. Figure 8B shows the responses of all voxels, across all subjects, as a two-dimensional histogram, aligned to the peak spatial frequency per voxel, plotted together with the model’s predictions. We can see the responses are symmetric about the peak, demonstrating that a log-Gaussian (as opposed to a linear Gaussian) function is the better choice. The responses do appear to deviate slightly from the model tuning curve: slightly flatter at the peak and falling faster away from it. A larger exponent could potentially improve the fit, e.g., exp(-log_2_(*x*)^4^) instead of exp(-log_2_(*x*)^2^). However, such a change will not have a large effect on the estimates of preferred spatial frequency, which is the primary focus of this paper.

To consolidate our findings, we combine the model parameters across subjects by bootstrapping a precision-weighted mean. For each parameter, we select 12 subjects with replacement, multiply each subject’s median parameter estimate by the precision of their response amplitudes (as estimated by GLMdenoise) averaged over all fit voxels, and average the resulting values. We then take this set of parameters and generate a set of predictions for the preferred period and gain across eccentricities and retinotopic angles, as well as for different stimulus classes (which determine the orientation seen by each voxel). We do this 100 times, plot the resulting median and 68% CI predictions in Figure 10, and plot the resulting median and 68% CI for the parameter values in Figure 9. We observe five distinct properties of the fitted functions:

**Figure 9:**
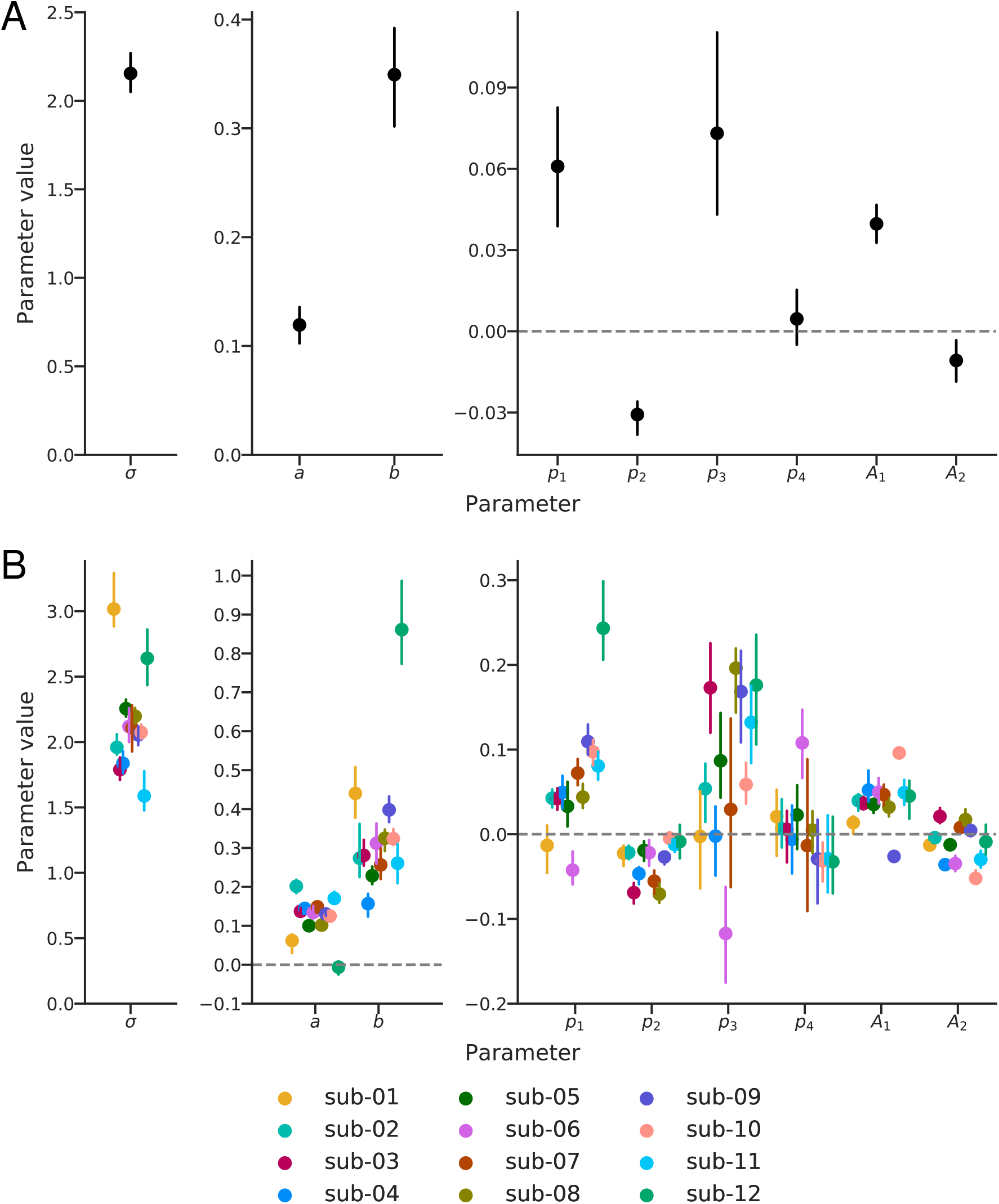
Parameter values (A) combined across all subjects and (B) in individual subjects. In both panels, median values ± 68% bootstrapped confidence intervals are plotted (note that *A*_3_ and *A*_4_ have been omitted, as determined from the previous model-selection analysis). (A) Parameter values obtained by bootstrapping parameter values across subjects from fits to the individual subject. A precision-weighted average is computed from each bootstrap. (B) Individual subject parameter values, bootstrapped across scans (as computed by GLMdenoise). A csv file containing these values (and instructions for use) can be found on the project GitHub.

**Figure 10:**
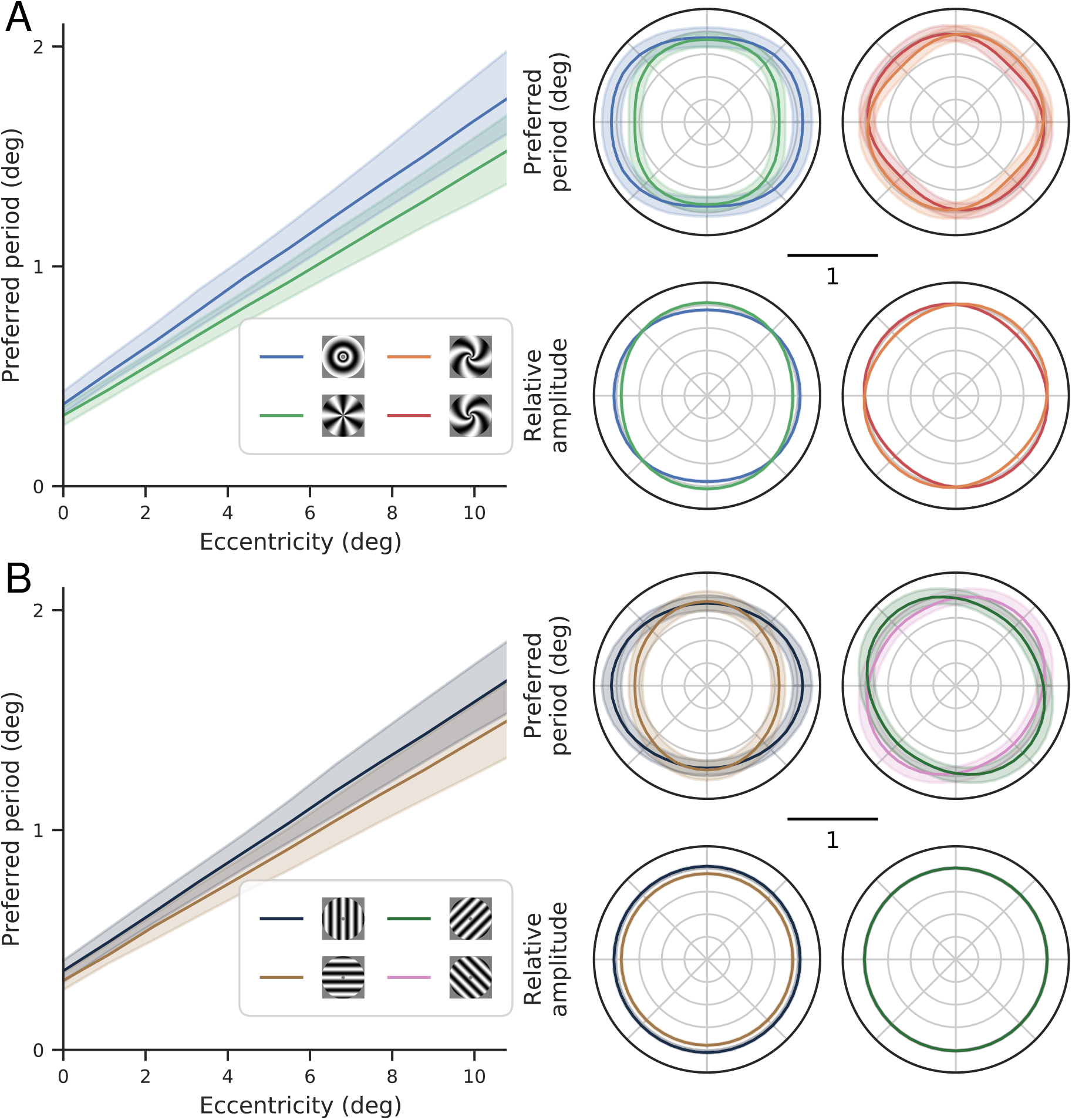
Spatial frequency preferences across the visual field in (A) relative and (B) absolute reference frames. In both panels, the left shows the preferred period as a function of eccentricity, top right shows the preferred period as a function of retinotopic angle at an eccentricity of 5 degrees, and bottom right shows the relative gain as a function of retinotopic angle (which does not depend on eccentricity; note that this relative gain does not change across voxels, only within a given voxel for different orientations). Only the extremal periods are shown in the left plot, for clarity (the others lie between the two plotted lines), and the cardinals and obliques are similarly plotted separately in the right plot for clarity. The predictions come from the model with parameter values shown in figure 9A, with the lines showing predictions from the median parameter and shaded region covering the 68% CI. Those parameters result from bootstrapping a precision-weighted average to combine the parameters from each subject’s individual fit with this model. Compare left plot in panel (A) to figure 6B.

##### Preferred period is an affine function of eccentricity

Specifically, the preferred period, as a function of eccentricity, is well approximated by a line with a significantly non-zero intercept. As discussed in the introduction, preferred period cannot decrease to zero at the fovea, since this would imply an infinite preferred spatial frequency. However, our stimuli do not include the region around the fovea, and thus our data do not constrain frequency tuning in that region. As such, the fitting procedure could potentially have arrived at an intercept of zero, supporting a “hinged line” model in which the preferred period decreases linearly toward the origin and levels out at some minimal value, as proposed in Freeman and Simoncelli, 2011.

##### Preferred period is largest for annular stimuli

As also seen in the 1D analysis (section 3.1), the annular stimuli have the highest preferred period at each eccentricity (figure 10A, left). Unlike in the 1D analysis, we can now see that the difference between the annuli and pinwheel stimuli varies as a function of retinotopic angle, with the largest difference at the horizontal meridian, decreasing to almost 0 by the vertical meridian (figure 10A, top right). At the horizontal meridian, the median preferred period is 1.06 for annuli and 0.80 for pinwheels. This difference as a function of stimulus angle is equivalent to about 2 degrees of eccentricity at a constant stimulus orientation.

##### Preferred period is largest for vertical stimuli

A similar pattern is seen for the model predictions for horizontal and vertical stimuli, in which there is an overall difference, modulated by retinotopic angle (figure 10B): The difference between their preferred periods reaches its maximal value at the horizontal meridian and decreases to almost 0 by the vertical meridian. This dependency between the preferred period effect and retinotopic angle comes from the combination of the vertical and annular biases: at the horizontal meridian, both go in the same direction (i.e., a vertical stimulus is an annular stimulus) and thus the gap in preferred period between vertical / annulus stimuli and horizontal / pinwheel stimuli is large. At the vertical meridian, on the other hand, they oppose each other (i.e., a vertical stimulus is a pinwheel stimulus), and, since the size of the two effects is roughly equal, the gap in preferred period between vertical / pinwheel stimuli and horizontal / annulus stimuli is zero.

##### Gain is largest for vertical stimuli

The effect of stimulus orientation on gain is smaller than the effect on preferred period, but more consistent across subjects. According to the model fits, vertical orientations evoke the largest BOLD signal (highest gain) and horizontal orientations the lowest. The two diagonal orientations are intermediate. The forward and reverse diagonal stimuli do not differ in gain because model 9 does not fit parameters A3 or A4, which would differentiate them. The gain for annuli and pinwheels varies as a function of retinotopic angle, based on where they align with the absolute orientation. Thus, the annuli have the highest gain on the horizontal meridian (where their absolute orientation is vertical), the pinwheels have the highest gain on the vertical meridian (where their absolute orientation is vertical), and the spirals have the highest gain on their respective diagonals.

##### Spatial frequency tuning is broad

By examining Figure 9A, we see that the standard deviation (*σ*) of our model is about 2.2 octaves, equivalent to a full-width half-max of 5.1 octaves. (The variability in the estimate comes from bootstrapping across subjects and across runs, not from variation across voxels or stimulus orientation, neither of which we modeled.) The 2.2 octave standard deviation of the tuning function is large relative to the variation in peak tuning across the V1 map. For example, the difference in preferred period between a foveal voxel (0 deg eccentricity, 0.35 deg period) and a 10 deg voxel (about 1.6 deg period) is equivalent to 1 standard deviation of the foveal voxel’s tuning function (2.2 octaves).

#### 3.2.3 Preferred period is (slightly) anti-correlated with V1 surface area

We observed substantial differences in preferred period across subjects. For example, at 6 deg eccentricity, preferred period ranges from 0.78 to 1.49 deg across our 12 subjects. A natural question is whether our measured preferred period is related to other functional or anatomical measures in V1. We motivated our initial scaling hypothesis by presenting the idea that the preferred spatial frequency may be a constant number of periods per population receptive field, and thus should drop as pRF size increases. Could the variability in pRF size across subjects account for the variability we see in preferred period? Estimated pRF size is far less reliable than pRF location, and so instead we compare preferred period to V1 surface area, which gives more robust estimates (Himmelberg et al., 2021; Lerma-Usabiaga et al., 2021). The results can be seen in appendix figure S1, comparing the preferred period at 6 degrees eccentricity with the total V1 surface area across participants. Both values span a range of 2:1, but they are essentially uncorrelated with each other (*R*^2^: median −3.42 × 10^-3^, 68% CI: [-2.84 × 10^-1^, 9.75 × 10^-2^]).

#### 3.2.4 Effect of retinotopic angle

To keep the model parameterization tractable, we excluded effects of retinotopic angle on preferred spatial frequency (except as mediated via relative stimulus orientation). To get a sense for whether retinotopic angle alone has additional explanatory power in our dataset, we fit model 3 (*p* = *ar_v_* + *b*, no effect of stimulus orientation and no modulation of gain) to the median BOLD response estimates on the quarters of the visual field around the two horizontal meridians (*θ_v_* ∈ [0, *π*/4] ∪ (3*π*/4,5*π*/4] ∪ (7*π*/4,2*π*]), and the quarters of the visual field around the two vertical meridians (*θ_v_* ∈ (*π*/4, 3*π*/4] ∪ (5*π*/4, 7*π*/4]). The bootstrapped average across subjects of the preferred period as a function of eccentricity for these two variants is shown in figure S2A. We can see that the model fit to voxels near the horizontal meridians has a higher preferred period near the fovea and a lower preferred period in the periphery, with the two meridian-only variants crossing at around 3 degrees. The error bars in that figure represent both the within-subject difference between the two variants and the between-subject differences in preferred period; figure S2B shows the difference between the two variants, calculated within subjects and then bootstrapped across them. This effect is clearly reliable across subjects, and the difference of approximately −.27 at 11 degrees is about 16% of the average preferred period there. Since our stimuli are balanced across relative stimulus orientations, this suggests that there is an effect of retinotopic angle alone on spatial frequency tuning, though further characterization is needed.

## 4 Discussion

We’ve used a set of log-polar grating stimuli to efficiently estimate spatial frequency preference in fMRI voxels of human V1. We quantified the effects of eccentricity, retinotopic angle, and stimulus orientation on voxel preferred period and response gain. As expected, the strongest relationship is the dependency on eccentricity: on average–across stimulus orientation, retinotopic angle, and subject–the preferred period is an affine function of eccentricity, which grows with a slope of about 0.12 degrees per degree of eccentricity and an intercept of about 0.35 degrees at the fovea. Preferred period is also modulated systematically by both stimulus orientation and retinotopic angle. Along the horizontal meridian, the increase in preferred period from horizontal to vertical stimuli (or, equivalently, from annular to pinwheel stimuli) is roughly equivalent to that seen when increasing eccentricity by 2 deg. On the vertical meridian, preferred periods of horizontal/vertical stimuli are indistinguishable. The response gain also exhibited small but systematic variations with stimulus orientation. Horizontal stimuli have an approximately 8% smaller response gain than vertical stimuli throughout the visual field.

### 4.1 Strengths

Our results are obtained using a multivariate, stimulus-referred model. Typically, stimulus-referred modeling of fMRI signals either fits each voxel independently (voxel-wise modeling) or fits average responses across regions. Voxel-wise modeling (e.g., Kay et al., 2008) has the flexibility of allowing researchers to place few or no constraints on the relationship of models across voxels. This flexibility comes with high parameter dimensionality: even a single visual area like V1, would typically require thousands of parameters, which can result in high noise sensitivity and lack of interpretability. Fitting models to regions of interest rather than voxels (e.g,. Boynton et al., 1996) reduces dimensionality, but loses cortical (and thus, retinotopic) resolution. Our method combines positive aspects of both approaches: it is sensitive to variability in the response properties across voxels, while placing constraints on how the parameters relate to each other across voxels to generate a useful and interpretable summary.

An advantage of the stimulus-referenced modeling approach is generalization. A 2D model of spatial frequency tuning is likely to simplify development of a more complete image-computable model of the visual cortex. Some image-computable models fit to fMRI or electrocorticography responses operate only on band-pass filtered images because they do not incorporate spatial frequency tuning (e.g., Hermes et al., 2019; Kay et al., 2013b; Kay and Yeatman, 2017). The stimuli were band-passed in these experiments to reduce the complexity of the image space. There have been some attempts to generalize models across scale but these have not been informed by a comprehensive set of measurements or models of spatial frequency tuning (Benson et al., 2017; Olman et al., 2017). A further advantage of the multivariate parametric approach is that it helps reduce bias from skewed voxel sampling. For example, there are fewer voxels near the vertical than near the horizontal meridian (Benson et al., 2020). In a voxel-wise fitting approach, preferences of voxels near the vertical meridian might be poorly fit or not fit at all (if no voxels have pRF centers along the meridian). Here, the parametric approach uses all the data to estimate each parameter, allowing better estimates for locations with limited data. Finally, a parametric model facilitates comparison across studies, as other measurements of spatial frequency tuning might not sample the identical orientations, spatial frequencies, and visual field locations.

### 4.2 Limitations

Our modeling approach has at least two important limitations. First, the characterization of the V1 maps is based on fMRI measurements, which combine the integration of the fMRI measurement (blood oxygenation within voxels) and the selectivity of those neurons linked to the changes in blood oxygenation. Some aspects of our results, such as the substantial additive offset at the fovea, may be particularly affected by these additional sources of integration. Comprehensive measures of spatial tuning across the entire map at the level of individual neurons do not exist. Models that explicitly account for both tuning of individual neurons and measurement pooling functions, such as Haak et al., 2012; Keliris et al., 2019 will be important for clarifying the relative contributions of these sources.

Second, our analysis assumes that for each voxel and each stimulus, there is a single spatial frequency and orientation driving the response. Because both stimulus properties varied continuously across our images, this use of the instantaneous frequency approximation is only valid locally. We think the effects are likely small since V1 receptive fields are relatively small and our stimulus properties varied gradually. In later stages of the visual system, where receptive fields are substantially larger, this use of instantaneous frequency will not well approximate the relevant portion of the stimulus for neuronal responses.

Finally, residual eye movements (microsaccades) could affect our results by increasing the positional uncertainty of the stimuli, or by effectively blurring them due to temporal integration. We think these effects are likely to be small (see appendix section 6.2 for more discussion), but we cannot entirely rule them out.

### 4.3 Related fMRI studies

A number of previous studies have reported spatial frequency preferences at multiple eccentricities in human V1 using fMRI. A comparison of those findings shows a wide range of estimates (figure 11; Aghajari et al., 2020; D’Souza et al., 2016; Farivar et al., 2017; Henriksson et al., 2008; Kay, 2011; Sasaki et al., 2001). With the exception of Aghajari et al., 2020 and Henriksson et al., 2008, these studies did not pursue the question of V1 spatial frequency tuning as their main question. All studies agree that preferred period should grow as an affine function of eccentricity, but the exact values for the slope and intercept vary widely. Overall, our results are most consistent with those of Aghajari et al., 2020. These studies fit tuning curves to different voxels or bands of voxels and plotted the peak as a function of eccentricity (sometimes, as in Aghajari et al., 2020, also separately plotting this for different quadrants of the visual field), similar to our 1D fits shown in figure 6A. The variability across studies could be due to many factors, including display calibration, analysis methods, temporal frequency of stimulus presentation, and the wide variety of spatial patterns used, from natural images in Kay, 2011 to phase-scrambled noise in Farivar et al., 2017 to plaids in Hess et al., 2009. Resolving the discrepancies may require use of multiple stimulus classes and analysis methods in the same study.

**Figure 11:**
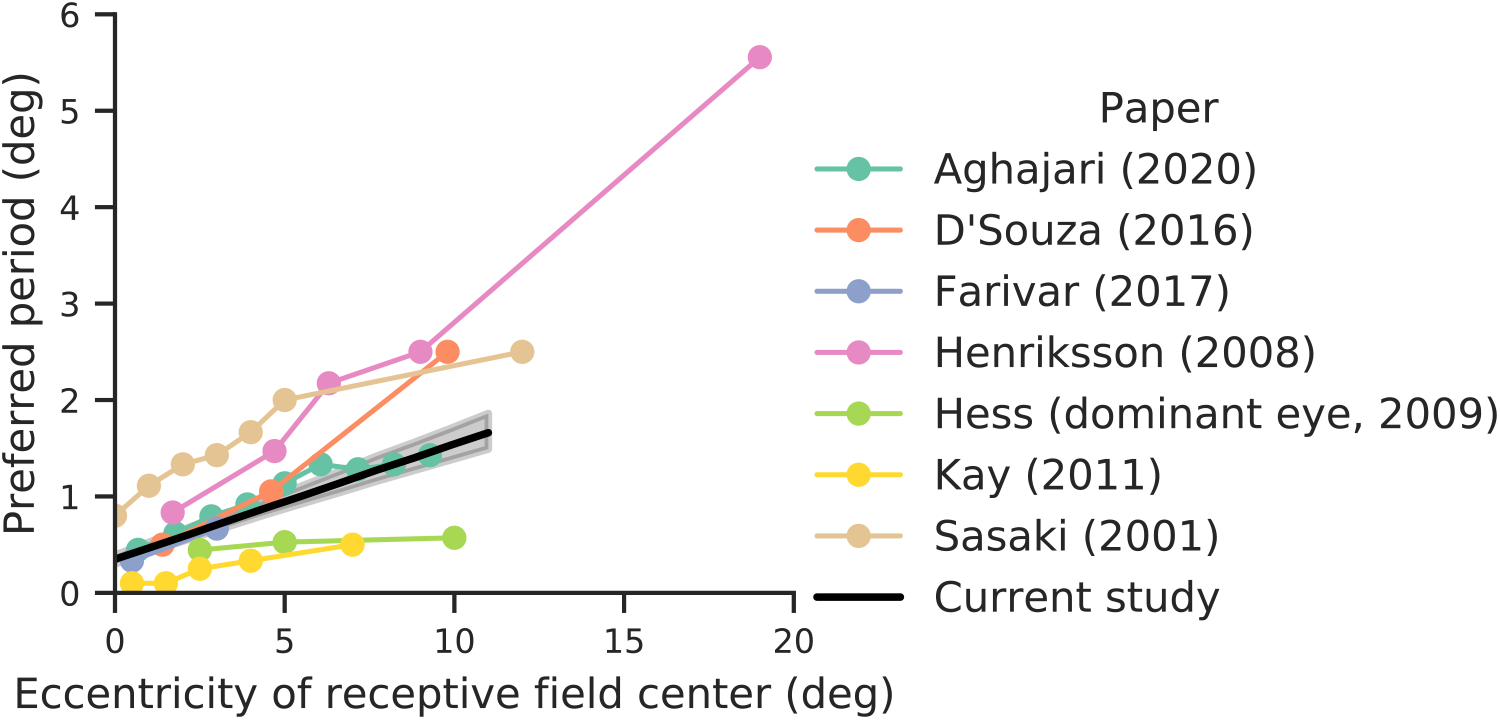
Comparison to previously reported eccentricity-dependence of spatial frequency measured with fMRI. All results show the preferred period at that eccentricity in V1 (all papers reported preferred spatial frequency; the reciprocal of that is shown). All values were estimated from published figures and are thus approximate. Black line represents our result, averaged across stimulus orientation and retinotopic angle, with line showing the median and shaded region the 68% confidence interval from precision-weighted bootstrap across subjects, as in figure 10.

Our 2D model assumes constant bandwidth in octaves. Aghajari et al., 2020 investigate the bandwidth of voxel spatial frequency tuning in more detail, concluding that it grows at a constant rate from approximately 3 octaves near the fovea to about 4.3 octaves at 9 degrees. Our model, like theirs, assumes a log-Gaussian tuning curve (figure 8B), but our bandwidth estimate of 5 octaves is larger than any of the values they observe in V1. We see no obvious explanation for the discrepancy. De Valois et al., 1982 measure spatial frequency bandwidth in macaque V1 simple and complex cells at multiple retinotopic locations and find a median bandwidth of approximately 1.5 octaves (similar across cell types and locations); they also show a negative correlation between peak spatial frequency tuning and spatial frequency bandwidth, with some low-pass neurons having a bandwidth of up to 3.25 octaves. Since neurons with a variety of spatial frequency tunings are found at any given retinotopic location, it is expected that V1 voxels would exhibit broader tuning than individual V1 neurons. This parallels findings in spatial receptive fields, which show larger sizes when measured for voxels with fMRI than measured in single units (Dumoulin and Wandell, 2008; Keliris et al., 2019).

### 4.4 Orientation tuning

There has been a long debate in the literature about whether orientation tuning is detectable in the BOLD signal on the spatial scale of voxels and, if so, what that means (Carlson, 2014; Freeman et al., 2013; Kamitani and Tong, 2005; Roth et al., 2018). Our model is recovering some degree of orientation tuning: with non-zero *A*_4_ and *A*_2_ values, response varies sinusoidally as a function of orientation. More specifically, we find an overall bias for vertical gratings. Freeman et al., 2013 found a mix of vertical bias near the fovea and a radial bias (e.g., voxels along the horizontal meridian preferring horizontal gratings) in the periphery. While our model agrees with the first finding, we find no evidence for the second (though our model does not allow for categorically distinct responses in the fovea and periphery, and fitting them separately may find some evidence for this, similar to the issue of retinotopic angle, see 3.2.4 and figure S2). However, we hesitate to interpret these results too strongly; as Carlson, 2014 and Roth et al., 2018 point out, orientation biases can be induced in the BOLD response by the stimulus presentation even with unbiased underlying neuronal responses. Further work is needed to tease out the source and implication of orientation tuning.

### 4.5 Scale and rotation invariance

An idealized model of visual system organization is that spatial frequency tuning (preferred period) is proportional to eccentricity, while being independent of polar angle and stimulus orientation. For example, the log-polar model of the warping of the visual field onto the V1 cortical surface by Schwartz, 1980 has these properties. The scaling with eccentricity has been proposed by Schwartz and others (Van Essen and Anderson, 1995) to endow the system with invariance to dilation and rotation (for transformations centered at the fovea), enabling percpetual generalization (but see Cavanagh, 1982 for a different interpretation). Our model fits show systematic deviations from each of these three properties.

First, we find that preferred period grows as an *affine* function of eccentricity, with a non-zero intercept. Independent of any measurements, one would not expect basic properties such as receptive field size to grow proportional to eccentricity due to limits at the fovea (the optics and cone apertures set upper bounds on resolution.) One simple correction to the idealized scaling model is adding an offset, or affine transform, as we find here. This is consistent with some models of cortical magnification in V1 (Benson and Winawer, 2018; Horton and Hoyt, 1991). An alternative model form is piece-wise linear (e.g., a “hinged line”), with proportional growth that flattens out at some minimal value near the fovea (as Freeman and Simoncelli, 2011 used in describing ventral stream receptive fields). This allows scale invariance outside the flat, foveal region. Our results show decisively that an offset is a better fit than a hinged line. The effect is relatively large: the offset at the fovea (preferred period of 0.35 deg) is equivalent to the difference in preferred period between 0 and 3 deg eccentricity. A substantial offset implies that the human V1 representation in the center of the visual field does not approximate a scaling rule, as also noted by Cavanagh, 1982. Given the importance of foveal vision for object recognition, the deviation from an idealized scaling rule at the fovea may have important implications for perception. Size judgments are in fact not invariant to eccentricity (Newsome, 1972) and have been shown to track individual differences in the topography of V1 (Moutsiana et al., 2016).

Second, we show spatial frequency tuning depends on orientation at the horizontal meridian, but not at the vertical meridian (see Figure 10, right panel). This is because the preferred period tuning for absolute orientation (vertical > horizontal) and for relative orientation (annuli > pinwheels) add for locations on the horizontal meridian, but cancel for locations at the vertical. In separate analyses, we also observed an overall higher peak spatial frequency for visual field quadrants near the horizontal meridian than the vertical outside of the central 3 deg, consistent with Aghajari et al., 2020. These results suggest that the quality of spatial representation will depend on polar angle. This is consistent with a large body of psychophysical results showing that performance on various tasks, including spatial resolution and contrast sensitivity, depend on stimulus polar angle, with better performance along the horizontal meridian than the vertical meridian and better performance along the lower vertical meridian than the upper vertical meridian (see Himmelberg et al., 2020 and the citations therein).

Finally, we show an overall annular bias in preferred spatial frequency: for any location in the visual field, an annular stimulus will have the lowest preferred spatial frequency, this bias varies across retinotopic angle, and increases with eccentricity. Few studies have examined the combination of stimulus orientation and retinotopic angle with sufficient resolution to determine whether an orientation effect is relative or absolute. An exception is Wilkinson et al., 2016, who used interference fringes to examine sinusoidal grating acuity changes across the visual field, and found that it is proportional to the sampling of retinal ganglion cells everywhere in the retina. Consistent with our study, they show that radial acuity is always higher than tangential acuity, that this effect is largest along the nasal horizontal meridian, and that the minimum angle of resolution (.5 / cutoff spatial frequency) grows roughly linearly with eccentricity. All told, this suggests that many, but not all, of the effects observed in the current study originate with the sampling of the midget retinal ganglion cell lattice.

## 5 Acknowledgments

The authors would like to thank Noah C. Benson for his assistance with the retinotopy analysis. They would also like to thank Shenglong Wang and the NYU HPC team for assistance in running this analysis on the NYU HPC cluster, as well as Shenglong Wang, Vicky Rampin, Rémi Rampin, Pablo Velasco, Deb Verhoff, and Kate Pechekhonova for assistance in sharing the data, code, and computational environment for this project. They would also like to acknowledge the following funding sources: NSF GRFP (WFB), HHMI (EPS), the Simons Foundation (EPS), NIH R01EY027401 (JW), and NIH R01EY027964 (JW).

## 6 Appendix

### 6.1 Stimulus properties

The local spatial frequency of our stimuli is equal to the magnitude of the gradient of the argument of cos(·) in Equation 1. Writing that argument as *g*(*r, θ*) = *ω_r_* ln(*r*) + *ω_a_θ* + *ϕ*, we differentiate to obtain the horizontal/vertical spatial frequency:

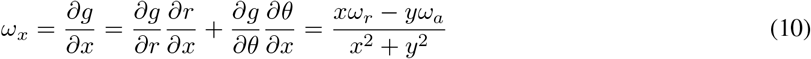

The local spatial frequency is then

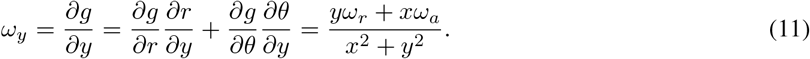

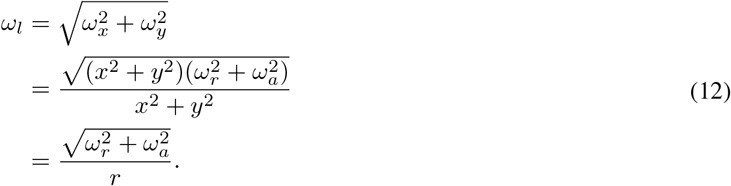

Thus, the local spatial frequency is proportional to the magnitude of the base frequency vector (*ω_α_, ω_r_*), decreasing as the inverse of eccentricity (r). To convert this from radians per pixel to cycles per degree, we multiply by a conversion factor, 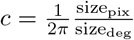. We use this measure of local spatial frequency when fitting tuning curves.

We can similarly find the local stimulus orientation, *θ_ι_*, by computing the angle of the frequency vector (*ω_x_, ω_y_*):

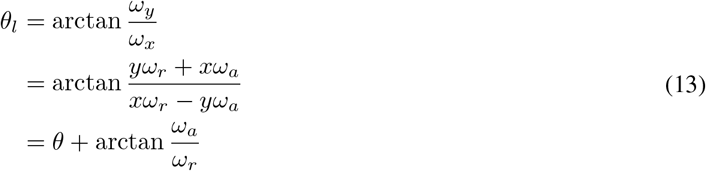

where 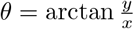. The local stimulus orientation is thus the sum of the angular location *θ* and the angle of the base frequency vector (*ω_r_, ω_α_*).

### 6.2 Behavior

Behavioral data is plotted in figures S3 and S4, combining across subjects and plotting them separately, respectively. When combining across subjects, there is no consistent pattern between performance and stimulus type: all stimulus types show similar behavior. When looking on a subject-by-subject basis, about half of the subjects show some differences across stimulus types. We might then worry that differences in eye movements or fixation stability may affect our results, such that any differences between spatial frequency tuning for annuli and pinwheels, say, are actually the result in differences in eye movements between those conditions. However, there appears to be no relationship between these behavioral patterns and the parameter values plotted in figure 9B. For example, sub-01 and sub-08 show similar behavioral patterns, with the highest miss rates for pinwheels, followed by forward spirals, then annuli and reverse spirals. However, their parameter values are not more similar to each other than to any other subject’s, making it unlikely that our parameter fits reflect differences in eye movement across stimuli.

Similarly, one might worry that differences in stimulus-independent eye movements might affect our results: fixational eye movements are known to be more common along the horizontal than vertical meridian, occuring at a rate of about 6 per minute (Thaler et al., 2013), with microsaccades in both directions having a median amplitude of 20 arcmin when there is a fixation target (Cherici et al., 2012). These increase the uncertainty of the spatial frequency within a voxel’s pRF, but, whenever voxels whose pRFs are on the right horizontal meridian see an increase in eccentricity due to a horizontal eye movement, voxels on the left horizontal meridian will see a decrease in eccentricity (and vice versa). Since our model does not allow left/right asymmetries in the fits, these two effects would approximately cancel.

**Figure S1:**
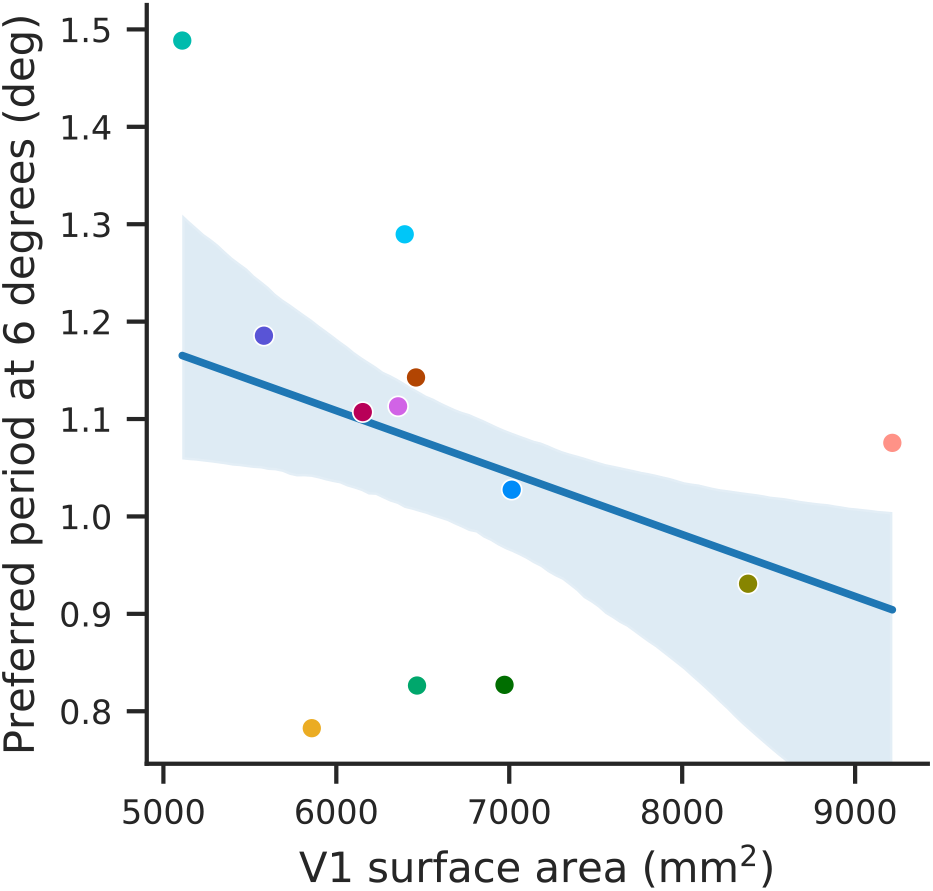
Scatter plot showing the preferred period at 6 degrees eccentricity against the total V1 surface area (both hemispheres) for all subjects. A middling eccentricity value was chosen so that the effects of both parameters a and b are visible. The two variables are essentially uncorrelated (line shows the median and the shaded region the 68% CI linear regression, bootstrapped across subjects). Subject colors are as in figure 9B.

Moreover, even if they did not fully cancel, microsaccades are relatively small compared to the observed effects of orientation (equivalent to about a 2 deg shift in eccentricity).

In addition to the uncertainty in voxel location discussed above, microsaccades combined with temporal averaging may blur the stimulus slightly, suppressing the high frequencies in our stimuli, which may shift our measured tuning curves to slightly lower frequencies. This effect would be most pronounced for those stimuli whose period is the same magnitude as the eye movements, which are present in our stimuli. This would increase the preferred periods on our plots and would have a larger effect on voxels at lower eccentricities. Eliminating or fully accounting for this effect is impossible given our setup, and future studies are necessary to account for its magnitude. Given the small size of fixational eye movements, we think their effects are likely to be small, but we cannot entirely rule them out.

### 6.3 Individual fits

Figures S5 through S11 show the individual subject fits for preferred period as a function of eccentricity from the 1d analysis (figure S5); preferred period as a function of eccentricity from the 2d model for relative (figure S6) and absolute (figure S7) reference frames; preferred period as a function of retinotopic angle (at 5 degrees eccentricity) for relative (figure S8) and absolute (figure S9) reference frames; and the relative gain as a function of retinotopic angle for relative (figure S10) and absolute (figure S11) reference frames. In all, we show the median and 68% confidence intervals obtained from bootstrapping across that subject’s fMRI runs.

Note that sub-12’s results are an outlier: their preferred period does not change as a function of eccentricity (also visible in the parameter plots in figure 9B; their a = 0). The noise in their GLMdenoise fits does not suggest any problems with the quality of this data, and the quality of their retinopic maps is also consistent with the other subjects. Therefore, they have been included in the analyses presented in this paper.

**Figure S2:**
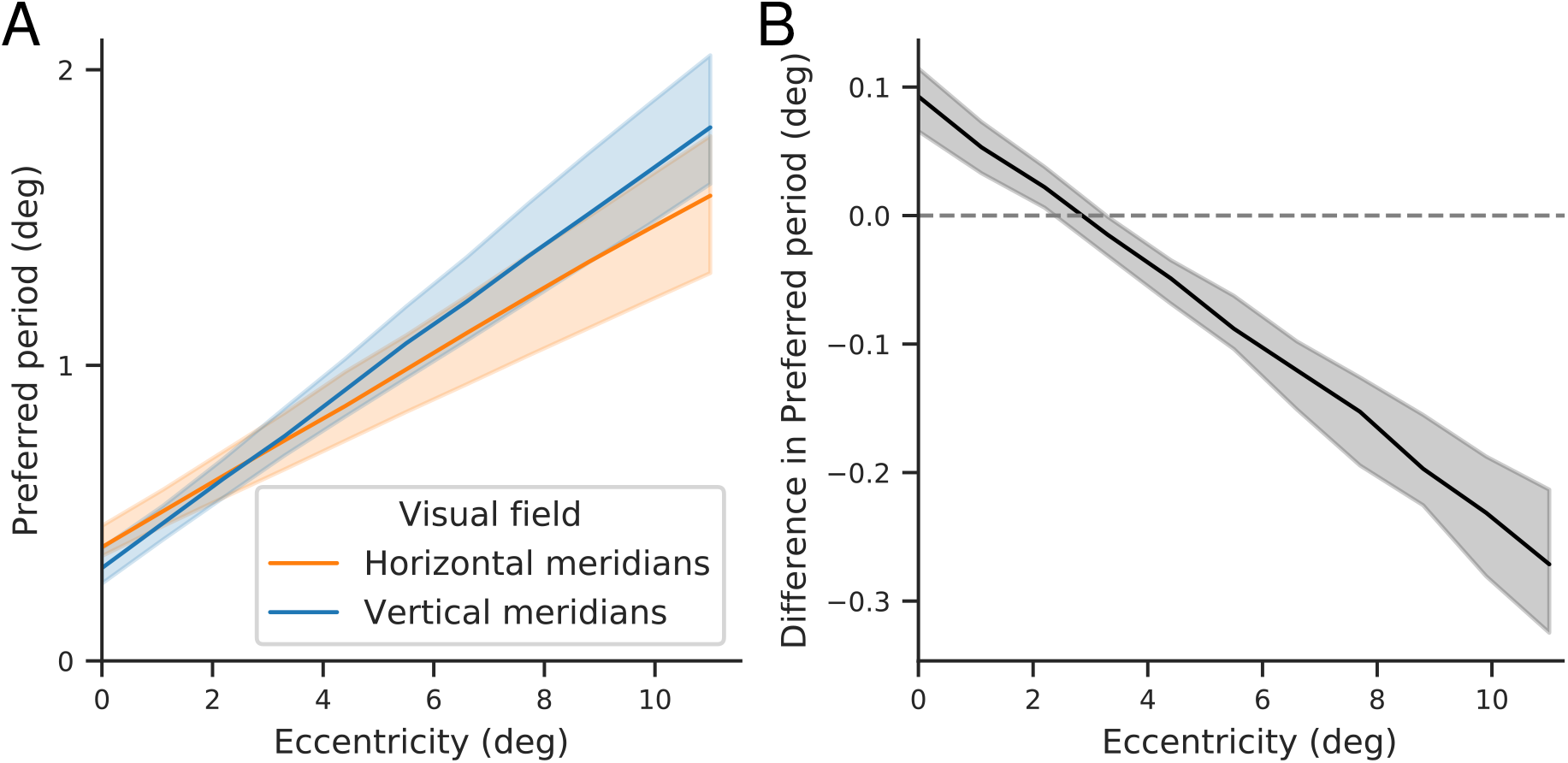
Voxels near the vertical meridians have a higher preferred period in the periphery and a lower preferred period near the fovea. Model 3 (*p* = *ar_v_* + *b*, no effect of stimulus orientation or modulation of gain) was fit to each subject’s median estimates of BOLD response. (A) Preferred period as a function of eccentricity for the two portions of the visual field. Lines and shaded region show the median and 68% CI from combining the preferred period across subjects by bootstrapping a precision-weighted average; uncertainty thus reflects both between-subject variance of preferred period and within-subject variance of the two visual field segments. (B) Difference between the preferred period in the voxels near the horizontal and vertical meridians, calculated within subjects, and then combined by bootstrapping a precision-weighted average.

**Figure S3:**
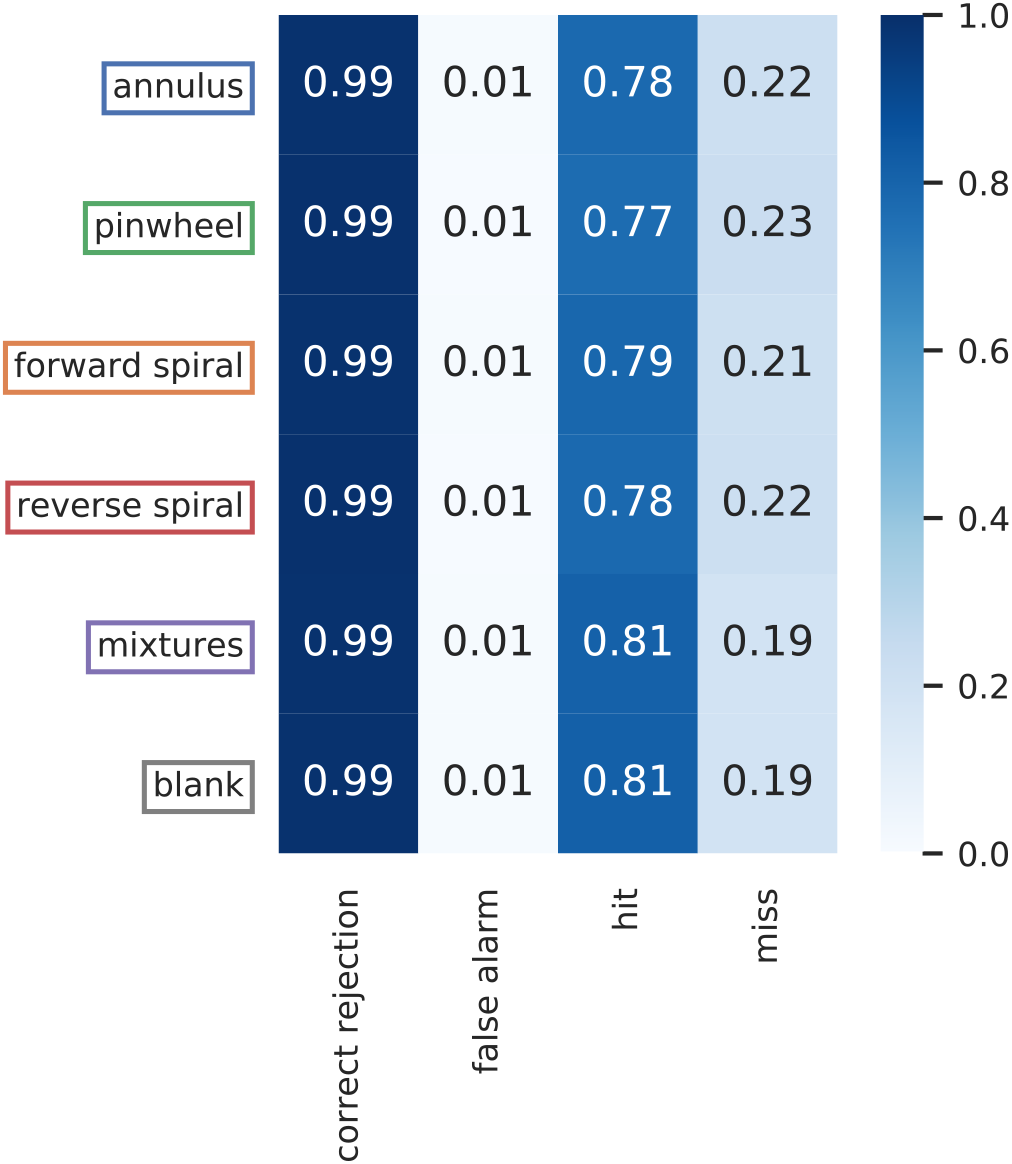
Summary of behavior combined across subjects. Stimulus type is indicated along the vertical axis (“blank” means there was no stimulus on the screen; these trials were interleaved throughout the scan as well as present at the very beginning and end), and outcome is indicated on the horizontal axis, with numbers giving the percentage of trials that fall into that category. Percentages and color are normalized so that the sum of correct rejections and false alarms is 1, as is the sum of hits and misses. During scans, subjects viewed a pseudo-random stream of digits at fixation and their task was to press a button whenever the digit repeated, which it did on one-sixth of the trials (the same digit was never shown three trials in a row). Behavior was consistent across stimulus types.

**Figure S4:**
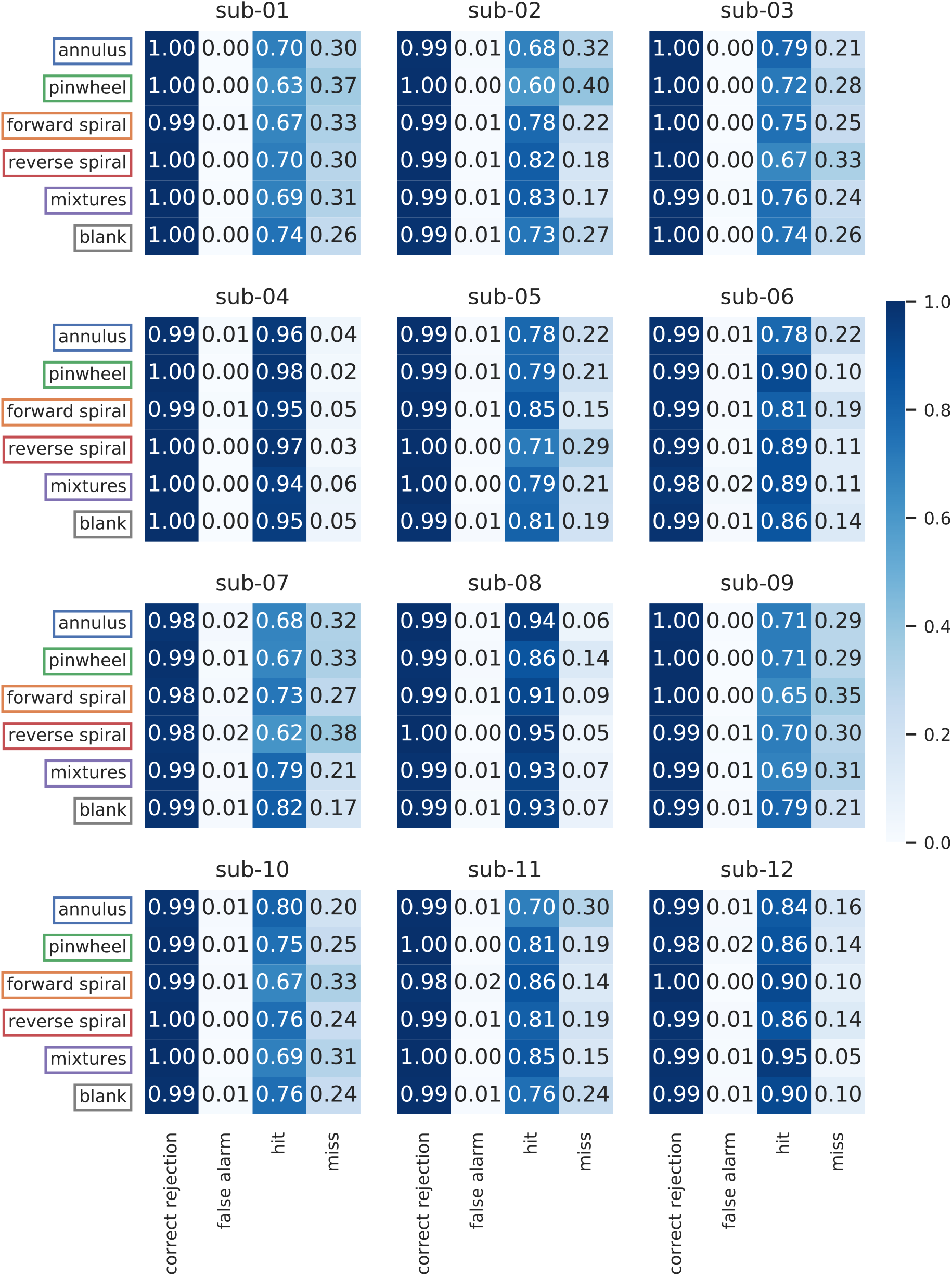
Summary of behavior on a per-subject basis. For details, see caption of S3. Performance varies across subjects (though false alarm rates are consistently low), but as in S3, there is no consistent difference in behavior across stimulus types.

**Figure S5:**
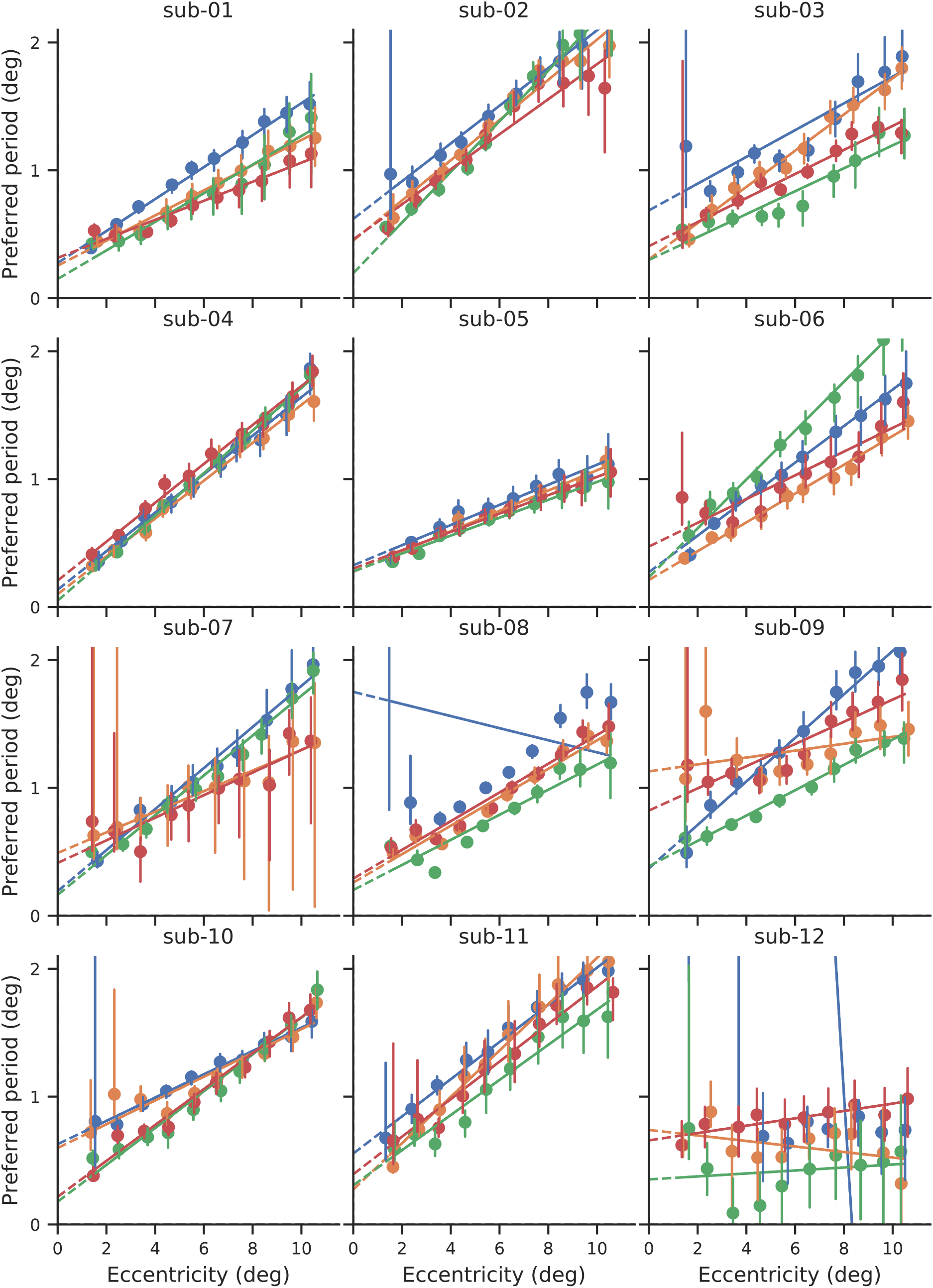
Individual subjects’ preferred period as function of eccentricity from 1d fits (as in figure 6A), for different stimulus classes. Points and vertical bars indicate the median and 68% confidence interval obtained from bootstrapping across fMRI runs.

**Figure S6:**
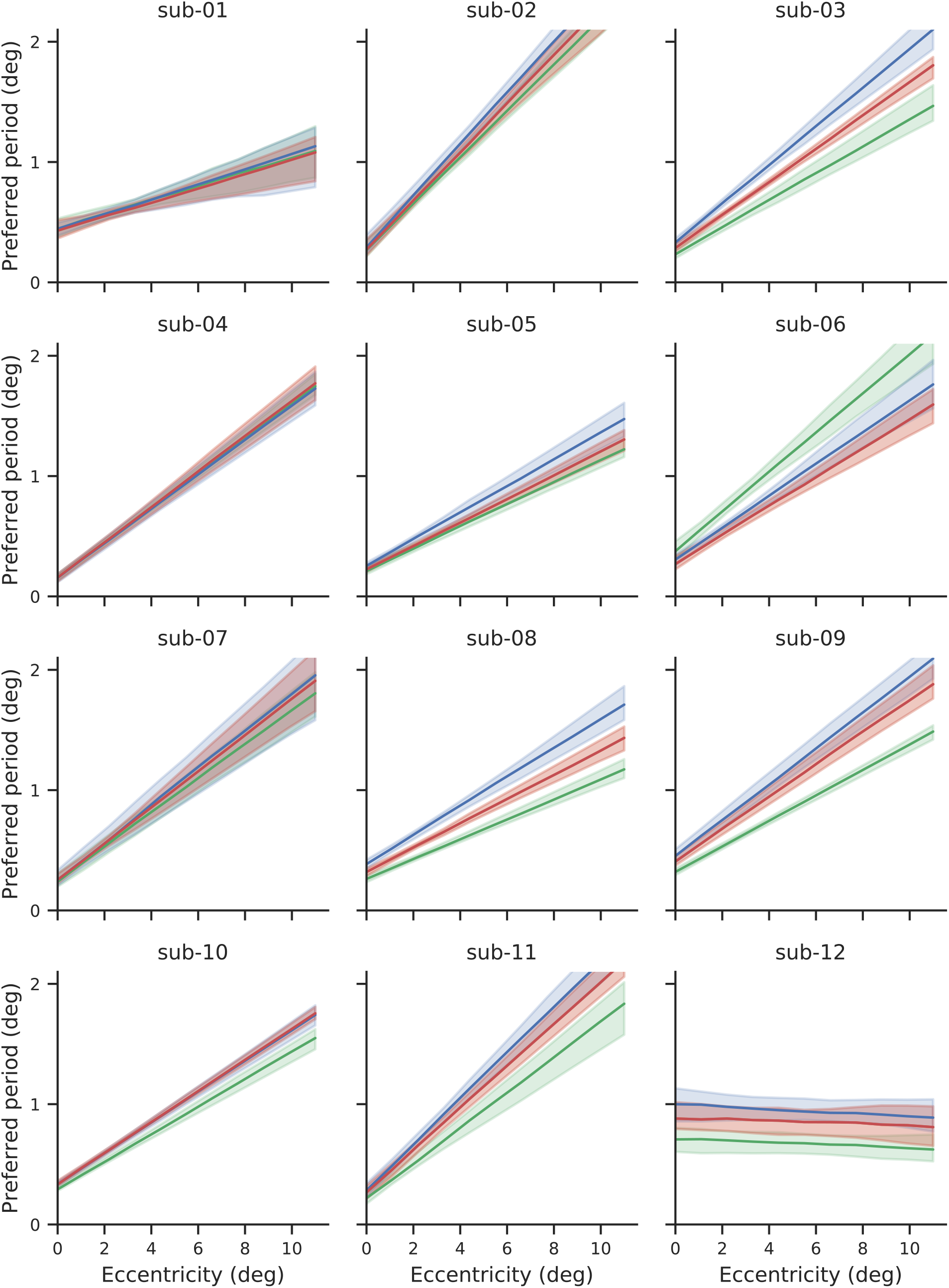
Individual subjects’ referred period as function of eccentricity from 2d model for relative reference frame (as in left panel of figure 10A). Averaged across all angles, lines show the median parameter and shaded regions cover the 68% confidence intervals obtained from bootstrapping across fMRI runs.

**Figure S7:**
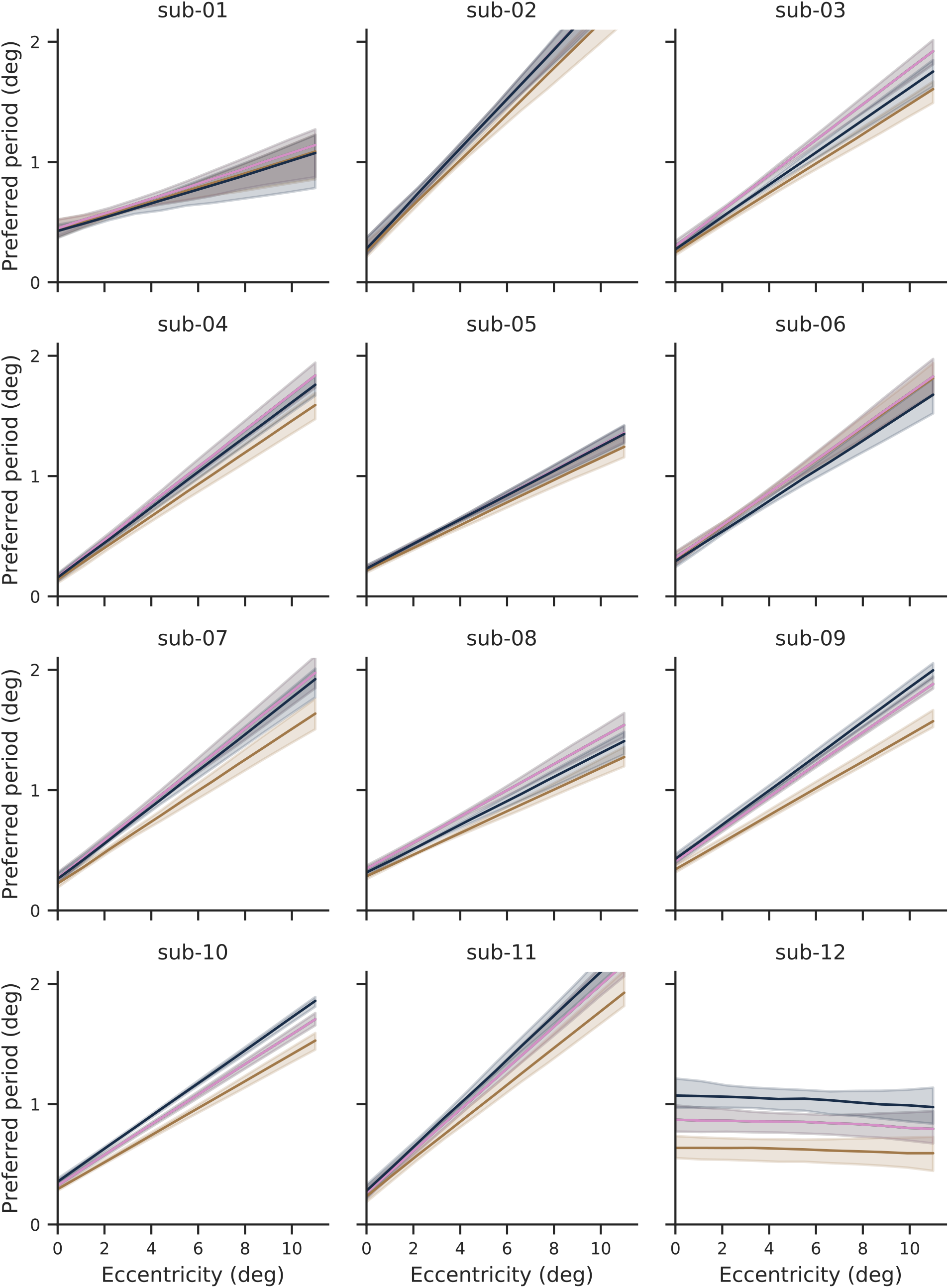
Individual subjects’ preferred period as function of eccentricity from 2d model for absolute reference frame (as in left panel of figure 10B). Averaged across all angles, lines show the median parameter and shaded regions cover the 68% confidence intervals obtained from bootstrapping across fMRI runs.

**Figure S8:**
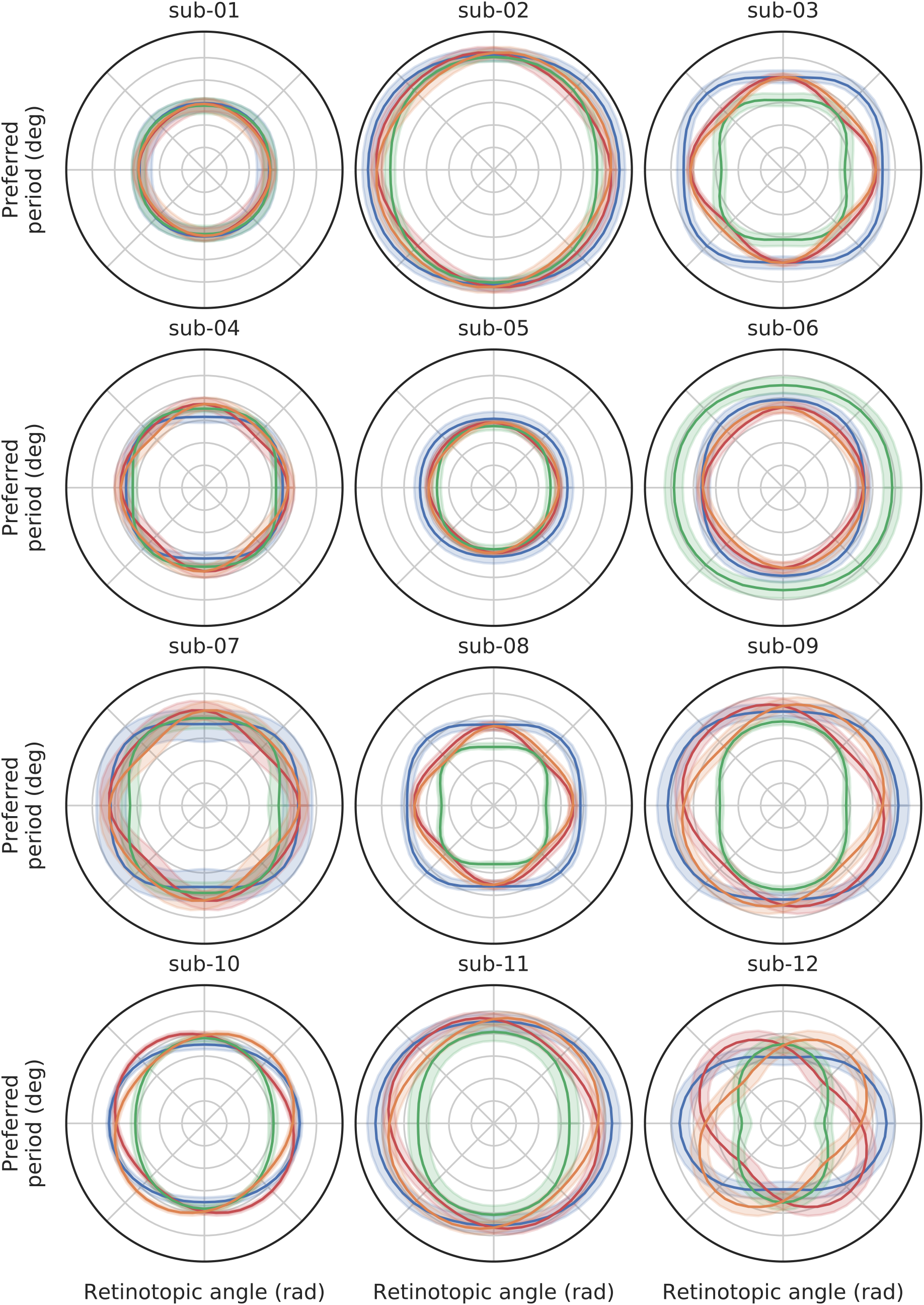
Individual subjects’ preferred period as a function of retinotopic angle at an eccentricity of 5 degrees for relative reference frame (as in top right panel of figure 10A). Lines show the median parameter and shaded regions cover the 68% confidence intervals obtained from bootstrapping across fMRI runs.

**Figure S9:**
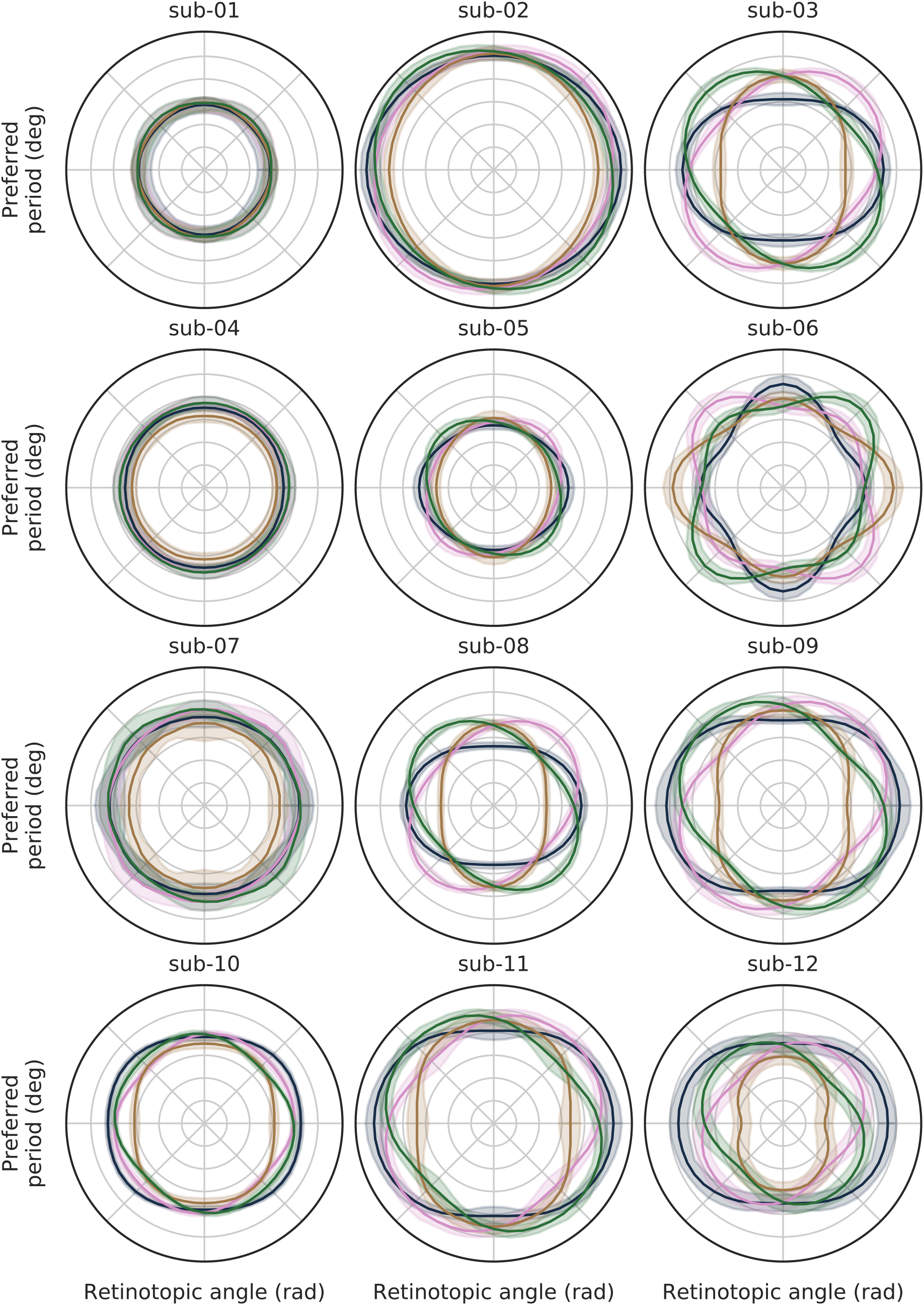
Individual subjects’ preferred period as a function of retinotopic angle at an eccentricity of 5 degrees for absolute reference frame (as in top right panel of figure 10B). Lines show the median parameter and shaded regions cover the 68% confidence intervals obtained from bootstrapping across fMRI runs.

**Figure S10:**
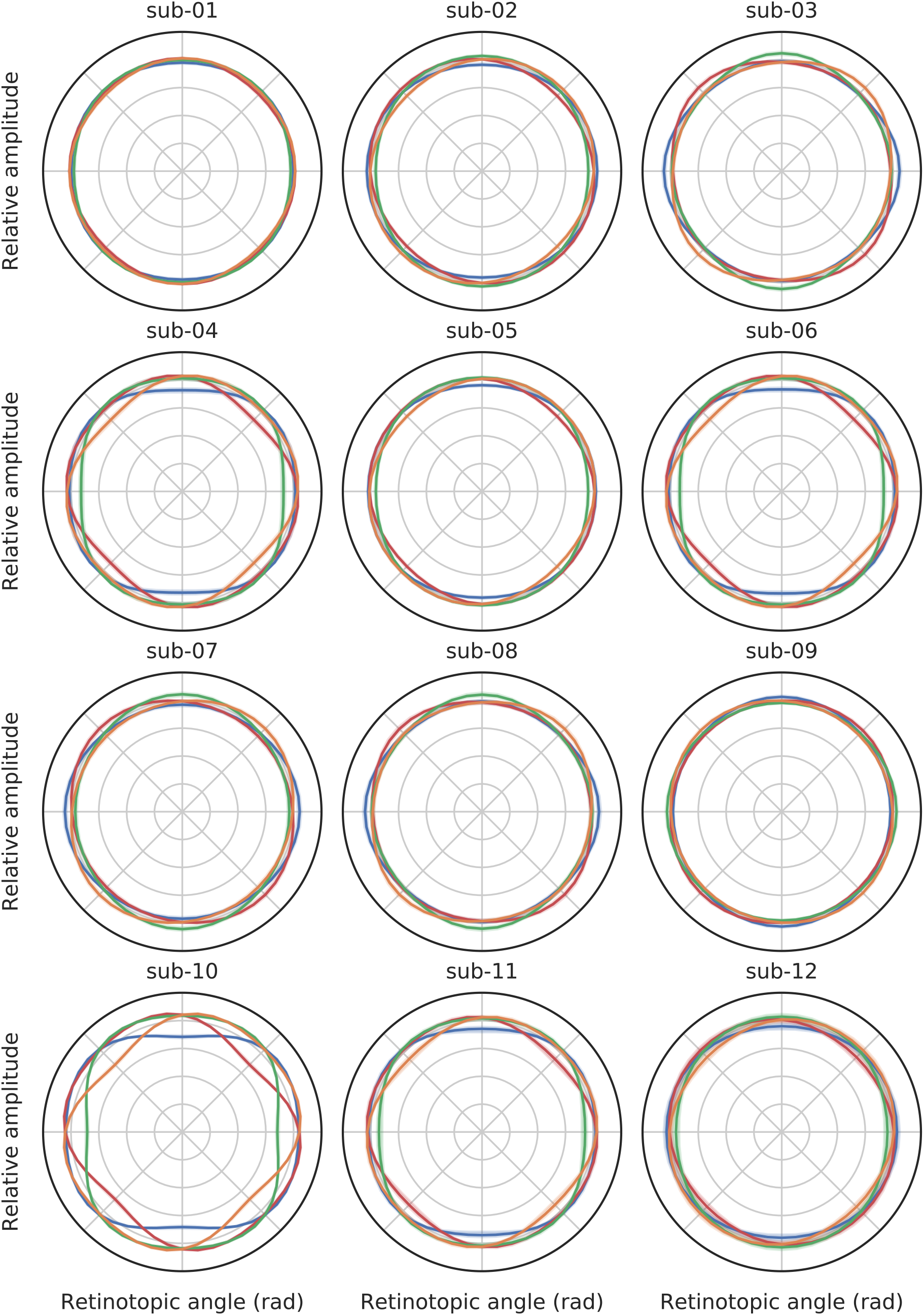
Individual subjects’ relative gain as a function of retinotopic angle for relative reference frame (as in bottom right panel of figure 10A). Lines show the median parameter and shaded regions cover the 68% confidence intervals obtained from bootstrapping across fMRI runs.

**Figure S11:**
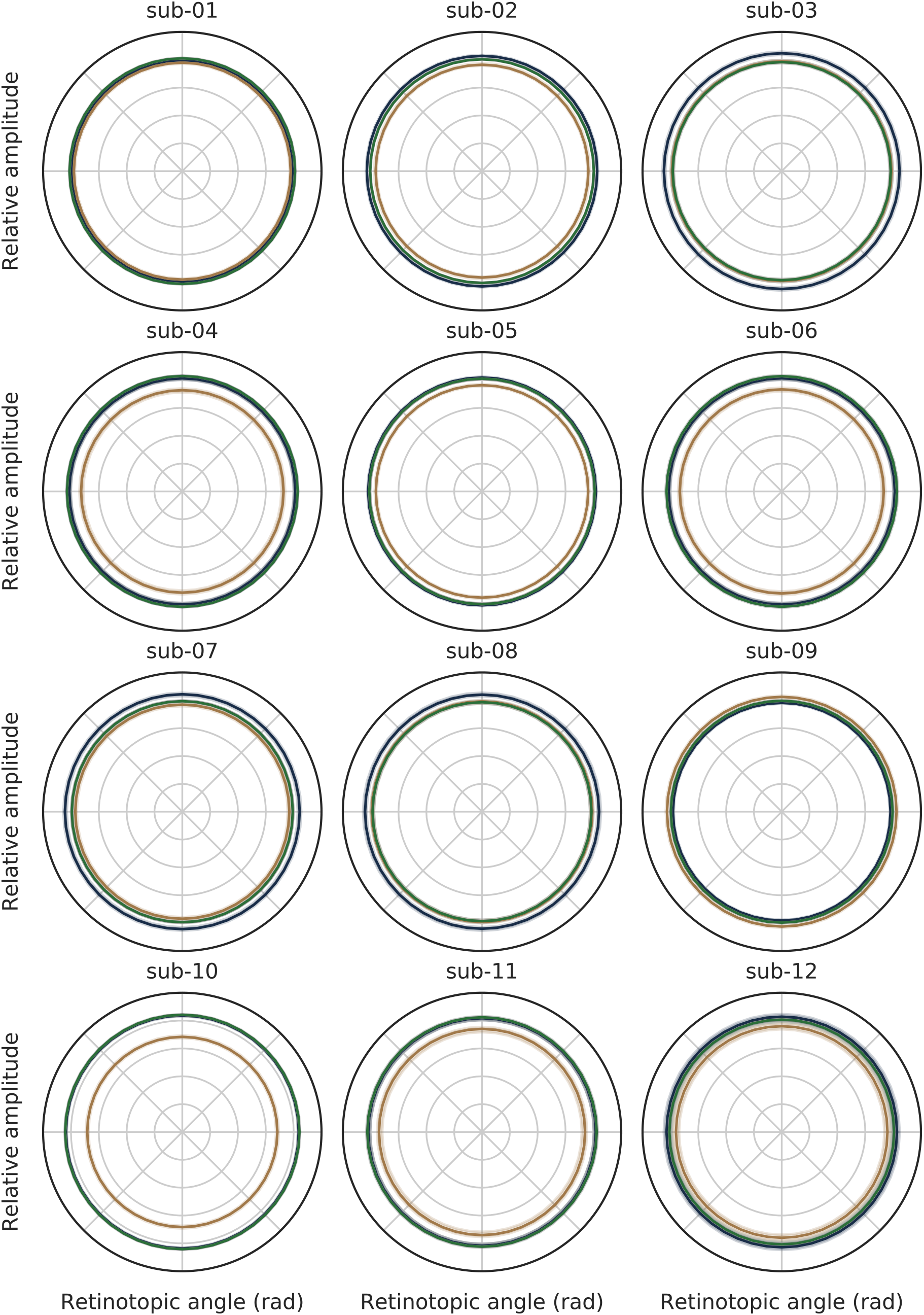
Individual subjects’ relative gain as a function of retinotopic angle for absolute reference frame (as in bottom right panel of figure 10B). Lines show the median parameter and shaded regions cover the 68% confidence intervals obtained from bootstrapping across fMRI runs.

## Footnotes

1 David Hubel described the process of characterizing visual field maps using single-unit electrophysiology as “a dismaying exercise in tedium, like trying to cut the back lawn with a pair of nail scissors” Hubel and Wiesel, 1977.

